# Dissecting oligogenic and polygenic indirect genetic effects through the lens of neighboring plant genotypes

**DOI:** 10.64898/2026.03.31.715746

**Authors:** Yasuhiro Sato, Kosuke Hamazaki

## Abstract

Individual phenotypes often depend on the genotypes of other individuals within a group. These phenomena are termed ‘indirect genetic effects’ (IGEs) and have been distinguished from direct genetic effects (DGEs) using quantitative genetic models. Recent studies have utilized high-resolution polymorphism data to enable genomic prediction (GP) and genome-wide association study (GWAS) of IGEs. To accelerate this practice, here we integrate polygenic and oligogenic IGEs using a multi-kernel mixed model. This model incorporates two random effects with a single covariance parameter for polygenic effects, while including neighbor allelic similarity to test SNP-marker effects for oligogenic effects. Our simulations demonstrated that the present model could infer a trade-off between DGEs and IGEs with a slight improvement in GWAS/GP performance. By applying our method to three species of woody plants, we found evidence for intergenotypic competition in aspen and apple trees, but limited evidence in climbing grapevines. Based on GWAS, we also detected significant variants associated with competitive IGEs on the apple trunk growth. Our study offers an alternative implementation for GWAS/GP of IGEs and thereby provides an effective tool to dissect the genetic architecture of mixing ability within a plant species.

## INTRODUCTION

Individual phenotypes often depend not only on an individual’s own genotype but also on the genotypes of other individuals within a group. This phenomenon, termed ‘indirect genetic effects’ (IGEs), is critical for evolutionary trajectories as it may alter the response to selection [1–3]. For instance, IGEs reduce trait heritability if they are negatively correlated with direct genetic effects (DGEs) [1,4]. In such a case, IGEs offer a valuable perspective for plant and animal breeding to optimize group performance by mitigating intraspecific competition [4– 6]. However, functional traits responsible for IGEs remain elusive [7]. For example, a set of crop traits designed to reduce competition, termed an ideotype [8], includes stem height, leaf angle, root length, and other dimensions of plant morphology [9,10]. To address the difficulty in identifying intermediate traits responsible for IGEs, many researchers have utilized genotype-based IGE models, which allow for trait-blind analyses of IGEs [1,5,7,11].

The application of genotype-based IGE models has been enhanced by the increasing availability of high-resolution and population-wide data of single nucleotide polymorphisms [3,12]. For instance, recent studies have conducted genome-wide association study (GWAS) and genomic prediction (GP) to dissect the genetic architecture of IGEs in humans [13,14], livestock [15], mice [7,16], *Arabidopsis thaliana* [11], and crops [17,18]. In laboratory mice, IGEs from cage mates have been analyzed for over 100 phenotypes and transcriptomes [7], where GWAS identified 24 quantitative trait loci (QTLs) underlying IGEs [16]. Besides mobile animals, IGEs have been analyzed for the model plant *A. thaliana* and crops to estimate mixing abilities among neighboring genotypes [17,19,20]. These pioneering studies have opened the way to implement GWAS and GP of IGEs.

In plants, IGEs occur among neighboring individuals because they cannot escape from their neighbors. These neighboring IGEs include multiple types of ecological interactions, such as competition and facilitation [21–24]. For instance, intraspecific competition can be shaped by a trade-off between individual and group performance, which corresponds to a negative DGE-IGE correlation [5,25]. Facilitation may also occur if resistant genotypes prevent pest spread on susceptible neighbors [20], which can shape a positive DGE-IGE correlation [5]. As a consequence of beneficial IGEs, a mixture of multiple genotypes sometimes yields more or less damaged than monocultures [19,20,26]. Such mixing abilities can be classified as general or specific mixing ability (GMA or SMA) [27–29]: The former indicates a single variety’s average ability irrespective to its counterparts, whereas the latter refers to specific combinations that interactively generate mixing ability. Enhancing mixing abilities by IGEs may therefore increase yield or resistance in agriculture [23,24].

To detect mixing abilities at the allelic level, our previous study developed a GWAS method named ‘Neighbor GWAS’ [30]. Specifically, Neighbor GWAS formulated frequency-dependent selection at each SNP locus to model the allelic SMA [12,31]. Furthermore, we needed to reduce false positive signals by correcting for the population structure shaped by additional factors such as IGEs [30]. To achieve this, we employed a two-step approach to reuse variance component estimates for fitting SNP-marker effects, which was similar to well-known GWAS algorithms such as EMMAX [32] and P3D [33]. By expanding this approach to a multi-kernel mixed model, we can implement Reliable Association Inference By Optimizing Weights called RAINBOW algorithm [34], thereby providing an efficient way to distinguish between DGEs and IGEs.

To enhance GWAS/GP of IGEs in plant species, here we provide an alternative model to integrate polygenic and oligogenic IGEs on a complex phenotype. We first presented a genotype-based IGE model by integrating Neighbor GWAS with RAINBOW. Subsequently, we conducted simulations to validate the performance of the present model against the original model. The present model was finally applied to three datasets of woody plant species. Based on the simulations and applications, we illustrated the feasibility of GWAS/GP of IGEs on a complex phenotype.

## RESULTS

### Model development

To enable flexible modeling of DGEs and IGEs, we developed a multi-kernel mixed model encompassing oligogenic and polygenic effects on a complex phenotype. These oligogenic and polygenic effects were modeled as fixed and random effects, respectively. The former enabled GWAS by testing each SNP, while the latter GP by estimating best linear unbiased predictors (BLUPs) based on genome-wide SNPs. We further mean oligogenic and polygenic IGEs by fixed and random effects from neighboring genotypes, respectively. To model neighbor genotypic interactions, let us assume that a focal individual *i* is interacting with neighboring individuals *j* in a local space *k*, which are designated as *k*_*i*_ and *k*_*j*_ (Fig. 1).

**Figure 1.**
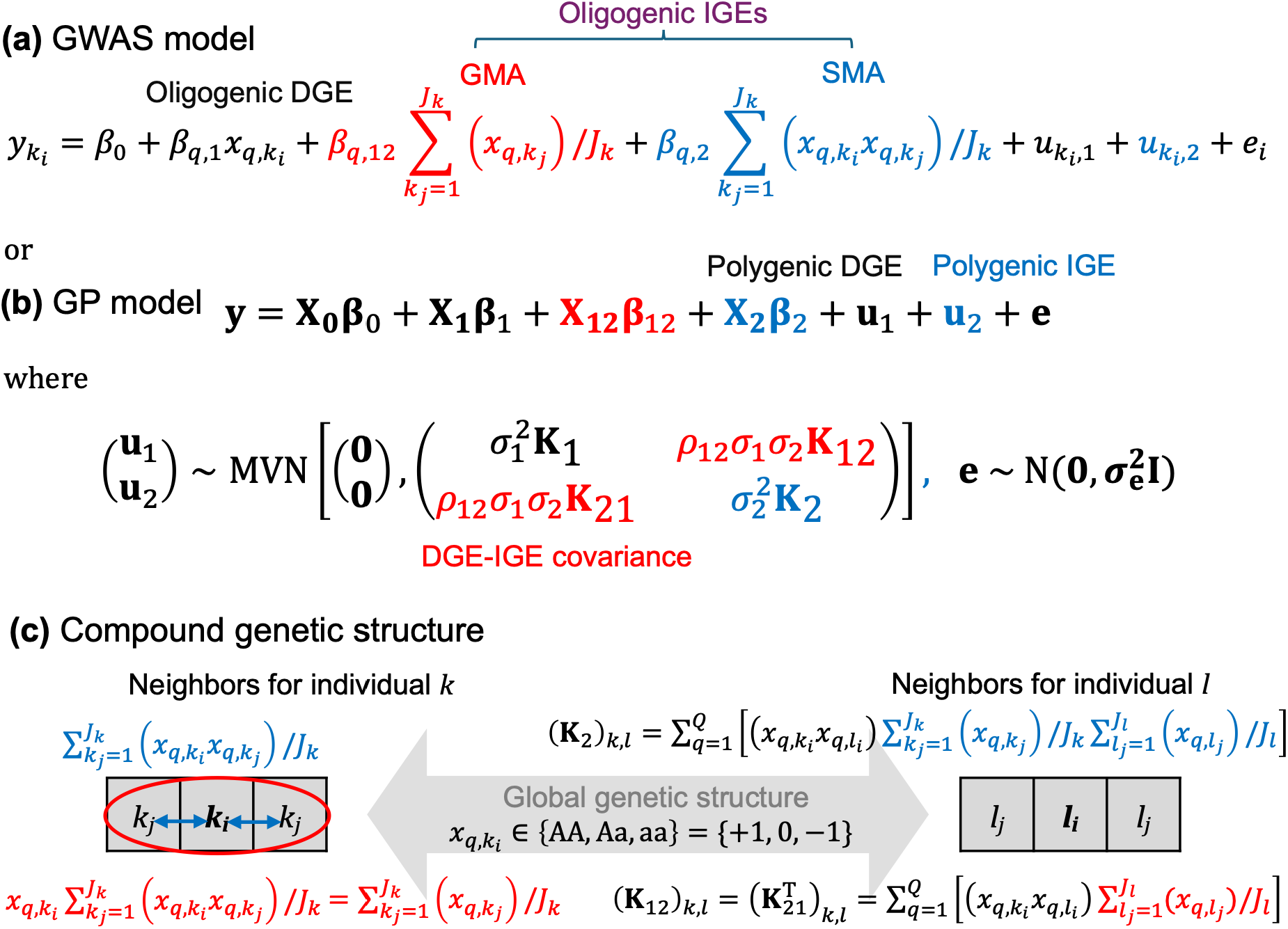
Overview of the proposed mixed model incorporating oligogenic and polygenic architectures of direct and indirect genetic effects (DGE and IGE). (a) Locus-level equation for genome-wide association study (GWAS). (b) Mixed model for genomic prediction (GP). Red and blue and red terms highlight IGEs related to general and specific mixing ability (GMA and SMA), respectively. Plain (black) terms indicate DGE components, which correspond to a standard GWAS/GP model. The model parameters are as follows; *x*_*q,i*_ ∈ {AA, Aa, aa} = {+1, 0, -1} whereby 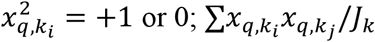: mean allelic similarity between focal and neighboring individuals; 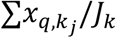: allele frequency bias near focal individuals; *β*_*q*,1_: SNP-wise DGE; *β*_*q*,2_: SNP-wise SMA of IGE based on neighbor allelic similarity; and *β*_*q*,12_: SNP-wise GMA of IGE derived from DGE×IGE based on the interaction term between *β*_*q*,1_ and *β*_*q*,2_. The random effects from polygenic DGEs **u**_1_ and IGEs **u**_2_ follow a multivariate normal (MVN) distribution with the DGE-IGE covariance *ρ*_12_. Panel (c) outlines how the weighted variance-covariance matrix incorporates the local (neighboring) and global spatial genetic structure for IGEs. The bottom right and left illustrations represent the neighboring space for individual *k* and *l*, which constitute the *k* and *l*-th elements of the four variance-covariance matrices as **K**_1_: Kinship matrix representing individual genetic similarity; **K**_2_: Kinship matrix weighted by neighboring allelic similarity; and **K**_12_ & **K**_21_: Kinship matrix weighted by neighboring allelic frequency, with the (co)variance-component parameters 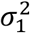: DGE genetic variance (>0); 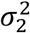: IGE genetic variance (>0); *ρ*_12_: DGE-IGE covariance (>0 or <0). See also “Model development” in the main text.

Following the original model of Neighbor GWAS [30], we then formulate IGEs of neighboring genotypes at *q*-th SNP locus on focal *k*_*i*_-th individual phenotype 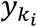 as follows.

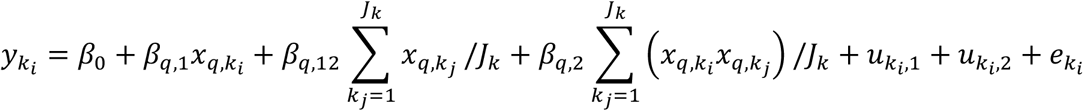

or

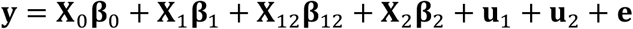

where

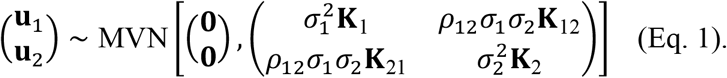

As for the fixed effects, 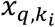 and 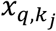 represent the genotype of focal *i*-th and neighbor *j*-th individuals in the local space *k* (where *J*_*k*_ is the total number of neighboring individuals). At the focal locus *q*, the two homozygotes are encoded as aa = -1 and AA = +1 with heterozygote as Aa = 0, i.e., 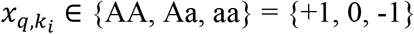. Accordingly, the fixed-effect coefficient *β*_*q*,2_ indicates the influence of mean allelic (di)similarity [i.e., (-1)×(+1) = (+1)×(- 1) = -1 or (+1)×(+1) = (-1)×(-1) = +1] between focal and all neighboring individuals on *y*_*k*_*i*. This formulation is analogous to the Ising model of ferromagnetics [35] when inter-genotypic interactions occur among neighboring individuals in a lattice space (e.g., sessile plants). In this analogy, *β*_*q*,2_ represents physical interactions among the south/north dipole of neighboring magnets while *β*_*q*,1_ corresponds to an external energy (Appendix S1; Fig. S1). In the context of evolutionary biology, the coefficient *β*_*q*,2_ represents positive or negative frequency-dependent selection when inter-genotypic interactions occur within a group [31] (Appendix S2 and S3; Fig. S2). The phenotype of one genotype depends on the others at a focal locus under frequency-dependent selection; therefore, we would designate frequency-dependent selection as a form of allelic level IGEs in this study. In addition to *β*_*q*,1_ and *β*_*q*,2_, *β*_*q*,12_ is defined as an interaction term between *β*_*q*,1_ and *β*_*q*,2_ because 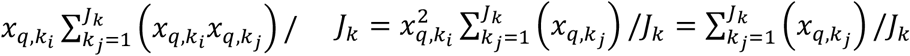 due to {AA, Aa, aa} = {+1, 0, -1} where 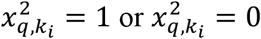 when 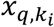 is heterozygous (Appendx S2). Theoretically, the sign of *β*_*q*,2_ determines whether the weighted mean of phenotypic values 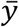 can be concave or convex against neighboring allele frequency (Fig. S2a and d), indicating SMA between two alleles at a focal locus. In contrast, *β*_*q*,12_ determines the slope of 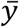 along the allele frequency (Fig. S2b-c and e-f) and thus indicates GMA of one allele irrespective of another. Although multiple equilibria become possible in the presence of heterozygote (Fig. S2b and e: cf. Fig. S2 of Sato et al. [31]), this property may not be relevant to cultivars for which we can manipulate genotype frequencies and their spatial arrangement. Therefore, the three coefficients *β*_*q*,1_, *β*_*q*,2_, *β*_*q*,12_ can be used as key parameters to distinguish between SNP-wise DGEs and IGEs in GWAS, which underpin the mean behavior of phenotypic values and consequent group performance under allelic mixture at a given locus. Hereafter we would refer oligogenic IGEs to both *β*_2_ or *β*_12_ but focus on *β*_2_ in the following GWASs to detect single-locus SMA of allelic mixture.

Besides the fixed effects, Eq. 1 includes random effects that can represent polygenic effects, i.e., BLUPs for DGEs and IGEs. These polygenic DGEs and IGEs are defined by **u**_1_ and **u**_2_, respectively. The fixed effects of Eq. 1 postulate individual-based intergenotypic interactions, whereby the design matrix should be an identity matrix (i.e., **Z** = **I**). In the multivariate normal (MVN) distribution, 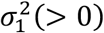 and 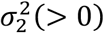 denote the genetic variance of DGEs and IGEs, respectively. The *N* × *N* kinship matrix **K**_1_ represents polygenic DGEs (where *N* means the total number of individuals), whereas one of the other variance-covariance matrices **K**_2_ represents polygenic IGEs. These two *N* × *N* variance-covariance matrices are defined by the cross-products 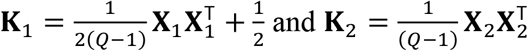 (where *Q* means the number of SNP loci). The *n*-th and *q*-th element of the SNP matrix **X**_1_ corresponds to the self-genotypic value as 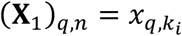. As we define 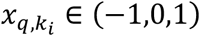, **K**_1_ is scaled so that its element represents the proportion of SNP loci shared among individuals. The *n*-th and *q*-th element of *N* individuals × *Q* SNPs matrix **X**_2_ is given as 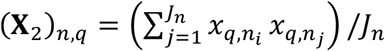. Accordingly, the *k*-th and *l*-th elements of *N* × *N* matrix **K**_2_ are given as 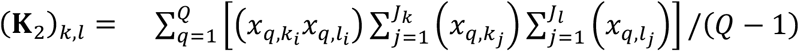. This indicates the polygenic effects (i.e., individual genetic similarity 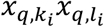) weighted by global allelic similarity between two local spaces *k* and *l* [i.e., a similarity metric of the mean allelic similarity in two local spaces 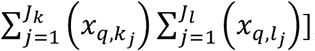, which is expected to correct for a spatial bias in allelic distribution in the GWAS of the neighbor genotypic effects *β*_*q*,2_ (Fig. 1b). Given also that plant genotypes are often randomized in a field space [20,36–39], we assume that residuals follow a uni-variate normal distribution as 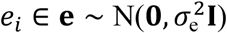 without covarying with IGEs.

When many loci underlie both DGE and IGE, these non-independent QTLs result in interactions between polygenic DGEs and IGEs (i.e., **X**_1_**β**_1_ and **X**_2_**β**_2_), thereby shaping covariance between **X**_1_**β**_1_ and **X**_2_**β**_2_ as Cov(**X**_1_**β**_1_, **X**_2_**β**_2_). Given that mean SNP effects are expected to be zero as *E*(**β**_1_) = *E*(**β**_2_) = 0, the DGE-IGE covariance is given as 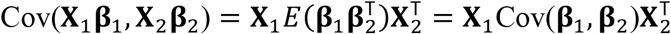 . When **β**_1_ and **β**_2_ are not independent, Cov(**β**_1_, **β**_2_) ≠ 0 and this covariance can be redefined as the MVN parameter

*ρ*_12_*σ*_1_*σ*_2_ = Cov(**β**_1_, **β**_2_). Meanwhile, 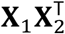 can be scaled and redefined as 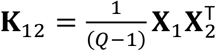 .The *k*-th and *l*-th elements of **K**_12_ are 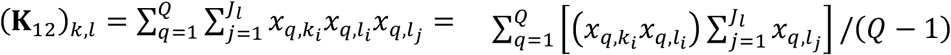, indicating the polygenic effects weighted by the local-space allele frequency in the neighborhood of focal individuals. Similarly, 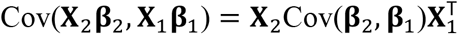 where 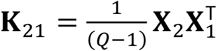. The *k*-th and *l*-th element of 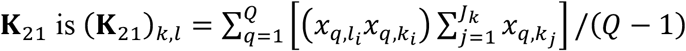. This indicates **K**_12_ ≠ **K**_21_, but 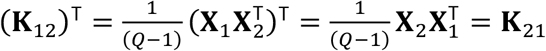. Furthermore, the local-space allele frequencies 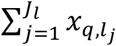 and 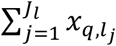 refer to the frequency of one allele at the same SNP locus. Thus, we can assume the same effect from the focal allele at a given locus as Cov(**β**_1_, **β**_2_) = Cov(**β**_2_, **β**_1_) = *ρ*_12_*σ*_1_*σ*_2_**I**. Combined with the two asymmetric variance-covariance matrices (i.e., 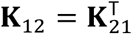), *ρ*_12_*σ*_1_*σ*_2_**K**_12_ and *ρ*_12_*σ*_1_*σ*_2_**K**_21_ represent asymmetric polygenic effects between focal and neighboring genotypes on 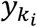. In GWAS, this random-effect covariance can be used to correct for population structure due to the locus-wise asymmetric neighbor effects *β*_*q*,12_.

In the case of Eq. 1, **u**_1_ and **u**_2_ are not independent due to their covariance. Therefore, the total phenotypic variance Var(**y**) can be decomposed as:

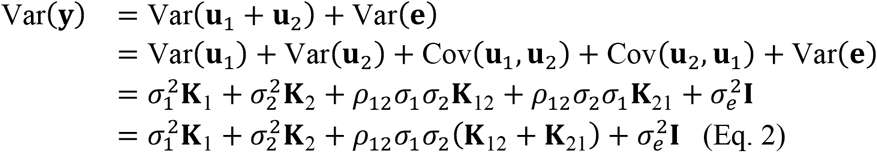

This Eq. 2 indicates that phenotypic variations can be partitioned into those explained by DGE, IGE, and their covariance in addition to residuals, where the three parameters take the range of 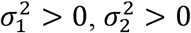, and *ρ*_12_ = (™∞, ∞). The sign of *ρ*_12_ therefore determines whether the DGE-IGE covariance increases or decreases phenotypic variance Var(**y**), which in turn affects response to selection.

We implemented the multi-kernel mixed model (Eq. 1) based on the RAINBOW algorithm [34]. This method works by optimizing the weights *w*_1_ + *w*_2_ = 1 for **K**_1_ and **K**_2_. In the present model, the variance component parameters are redefined as 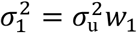 and 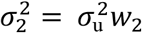, where 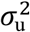 represents the total genetic variance due to the two random effects. Accordingly, the proportion of phenotypic variation explained (PVE) by DGE, IGE, and their covariance can be partitioned into PVE_DGE_ = *w*_1_PVE_u_, PVE_IGE_ = *w*_2_PVE_u_, and 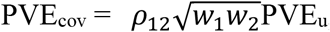, where 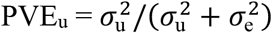. In this respect, PVE_cov_ is scaled by its kernel weight 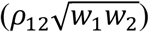 and the fraction of total genetic variance (PVE_u_), ranging from -1 to +1. Therefore, PVE_cov_ represents the sign and magnitude of PVE increased or decreased by the DGE-IGE covariance. Estimating variance component parameters yields a weighted kernel as 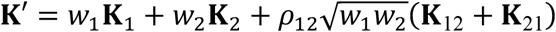. This weighted kernel allows us to perform GWAS by comparing models with or without each SNP-wise DGEs or IGEs for *β*_*q*,1_, *β*_*q*,2_, or *β*_*q*,12_ based on likelihood ratio tests.

### Benchmark simulation

Using simulated genomes and phenotypes, we tested the performance of the present model in variation partitioning, GP, and GWAS. To first address how correctly variance component parameters can be estimated, we performed independent iterations 30 times with varying covariance and ratio between DGE and IGE (Fig. 2; Fig. S4 and S5). These simulations showed that the mean of the estimated covariance parameter 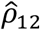 matched those of true models up to 10^-2^ order when DGEs and IGEs evenly contributed with a strong covariance (*s*_1_: *w*_2_ = 0.5: 0.5 with *ρ*_12_ ± 0.6: Fig. 2a and g). Even when we simulated moderate covariance, 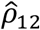 ranged over the true parameters within its SD (Mean ± SD = -0.38 ± 0.22 and 0.30 ± 0.07 for the true *ρ*_12_ = -0.3 and 0.3, respectively: Fig. S4e and i). When DGEs contributed far larger than IGEs (*w*_1_: *w*_2_ = 0.9: 0.1), 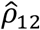 was slightly overestimated yet did not range across zero (Fig. S5a and h). The null scenario in which the true *ρ*_12_ = 0 confirmed that 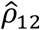 overlapped zero (Mean ± SD = -0.03 ± 0.08: Fig. S4a). These results demonstrate the capability of the present model in estimating the sign of DGE-IGE covariance.

**Figure 2.**
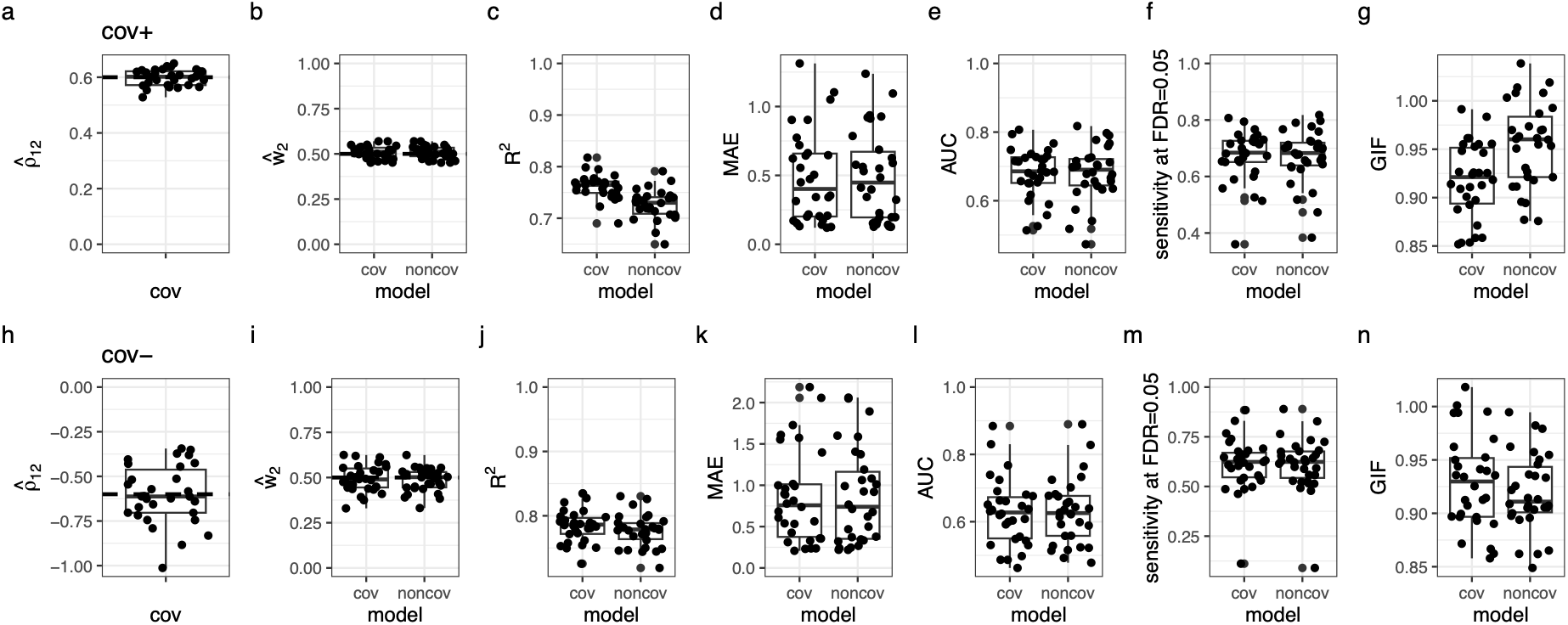
Benchmark simulations under the scenarios of positive or negative covariance with even DGE-IGE weights as *w*_1_: *w*_2_ = 0.5: 0.5. The upper panels (a-g) show the results of positive covariance (*ρ* = 0.6) while the lower panels (h-n) show those of negative covariance (*ρ* = ™0.6). (a and h) Estimated covariance 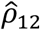 between polygenic DGE and IGE. Horizontal dashed lines indicate the true *ρ*_12_ value. (b and i) Estimated weight of IGE 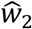 relative to that of DGE 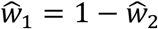. Models with or without the covariance 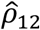 (cov or noncov) were fitted to simulated phenotypes. Horizontal dashed lines indicate the true *w*_2_ value. (c-d and j-k) Genomic prediction based on the models with or without the covariance 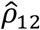 (cov or noncov). The model performance was evaluated using the coefficient of determination (*R*^2^: c and j) and mean absolute error (MAE: d and k) between the predicted phenotypic values **ŷ** and the observed phenotypic values of the test dataset **y.** (e-g and l-n) GWAS performance for the models with or without the covariance 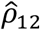 (cov or noncov) to detect causative IGE SNPs. The model performance was evaluated using the area under the ROC curve (AUC: e and l) and sensitivity at the stringent threshold at false discovery rate (FDR) = 0.05 (f and m). The genomic inflation factors (GIF: g and n) indicate the inflation (>1.0) or deflation (<1.0) of *p*-values compared to random expectation. Box plots display the median and quartiles among 30 independent iterations, with whiskers extending to 1.5 × inter-quartiles.

The relative importance of DGEs and IGEs was determined by the weight parameters *w*_1_ and *w*_2_ (where *w*_1_ + *w*_2_ = 1). By focusing on *w*_2_, we found that models with and without the covariance could both agree with the true weight when their contributions were even (i.e., *w*_1_: *w*_2_ = 0.5: 0.5: Fig. 2b and i; Fig. S4f and j). When true weights were biased to have small IGEs as *w*_1_: *w*_2_ = 0.9: 0.1, however, the model without *ρ*_12_ overestimated IGE weight 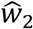 when true *ρ*_12_ was negative (Fig. S5i). This overestimation could be avoided by considering *ρ*_12_ (Fig. S5i). Regardless of the inclusion of *ρ*_12_, the overestimation of 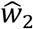 was not observed when true *ρ*_12_ was positive (Fig. S5b). Therefore, the models with the DGE-IGE covariance *ρ*_12_ can outperform those without the covariance especially when IGEs have far smaller contributions than DGEs.

Regarding the estimated range of (co)variance parameters, the scenarios of positive DGE-IGE covariance exhibited smaller variations in 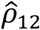 and 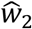 than those of negative DGE-IGE covariance (Fig. 2a-b and h-i). To further examine the parameter variability, we compared the simulated estimates with theoretically expected SEs that were calculated from the inverse of the expected Fisher’s information matrix (Appendix S5; Fig. S6). These SEs were larger when *ρ*_12_ < 0 than that of *ρ*_12_ > 0 (mean ± 95% CI of SE of 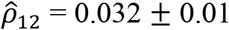 and 0.157 ± 0.05 for true *ρ*_12_ >0 and <0, respectively: Fig. S6a and b), providing the consistent range for simulated estimates (SD of 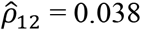 and 0.167for true *ρ*_12_ >0 and <0, respectively: Fig. S6a and b). Consistent with the results of 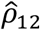, the SEs of the IGE weight were larger when true *ρ*_12_ was negative (Fig. S6c and d). The null scenarios in which the true *ρ*_12_ = 0 exhibited an intermediate level of variation in 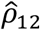 and 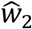, providing quantitative evidence between the cases of true *ρ*_12_ >0 and <0. These results suggest that, compared to the positive sign, the negative DGE-IGE covariance is relatively difficult for our model to detect.

To test whether the inclusion of the DGE-IGE covariance *ρ*_12_ could improve the performance of GP, we then calculated the coefficient of determination (*R*^2^) and mean absolute error (MAE) between predicted and simulated BLUPs. These two metrics were obtained by leaving the one-tenth of the simulated data as a test dataset, while the rest was a training dataset (Fig. 2c-d and j-k; Fig. S4c-d, g-h and k-j; Fig. S5c-d and j-k). This cross-validation procedure showed that the models with DGE-IGE covariance *ρ*_12_ slightly improved *R*^2^ compared to those without the covariance (mean of difference = 0.035 and 0.007, *t*_29_ = 22.5 and 15.8, *p* < 1.0^-15^ and 1.0^-15^ by a paired *t*-test for *ρ*_12_ = 0.6 and ™0.6, respectively: Fig. 2c and j). This pattern was also observed for the case of weak covariance (*ρ*_12_ = ±0.3: Fig. S4g and k) or small IGE variance (*w*_1_: *w*_2_ = 0.9: 0.1: Fig. S5c and j), although the extent of improvement was small especially when true *ρ*_12_ was negative. This small improvement in *R*^2^ occurred partly due to the approximation of the weighted variance-covariance matrix to its nearest positive definite matrix when *ρ*_12_ < 0 (Fig. S7). In contrast to the cases of true *ρ*_12_ ≠ 0, the models with *ρ*_12_ had slightly worse *R*^2^ when true *ρ*_12_ = 0 (paired *t*-test, mean of difference = ™0.0003, *t*_29_ = ™2.65, *p* = 0.013: Fig. S4c), indicating that inclusion of ineffective parameters decreases the model predictability. These results illustrate the power of the present model to improve GP when IGEs covary with DGEs.

We then tested whether the inclusion of the DGE-IGE covariance *ρ*_12_ could improve the performance of GWAS to detect causative IGE SNPs (i.e., non-zero *β*_*q*,2_). Under the distinct scenarios of positive or negative covariance *ρ*_12_ = ±0.6, we evaluated the model performance using the receiver operating characteristic curve (ROC) and its area under the curve (AUC). These GWAS simulations showed moderate power to detect causative SNPs for both cases of negative and positive covariance (median AUC > 0.6 and AUC < 0.8: Fig. 2e and l). This moderate power was also supported by the sensitivity at the stringent specificity (at FDR < 0.05: Fig. 2f and m) and remained even when IGEs were far less influential than DGEs (*w*_1_: *w*_2_ = 0.9: 0.1: Fig. S5e-l and f-m). These two metrics confirmed the feasibility of GWAS of IGE but its performance did not differ considerably regardless of the DGE-IGE covariance (Fig. 2e-f and k-m). Furthermore, we also calculated genomic inflation factors (GIFs) [40,41] to inspect the risk of false positive signals. We found no strong signals of *p*-value inflation (median GIF < 1.2: Fig. 2g and n), but GIFs exhibited contrasting patterns between the scenarios of true *ρ*_12_ < 0 and *ρ*_12_ > 0. The models with DGE-IGE covariance *ρ*_12_ decreased GIFs when true *ρ*_12_ > 0 (Fig. 2g; Fig. S5g), whereas they increased GIFs when true *ρ*_12_ < 0 (Fig. 2n; Fig. S5n). These results suggest that both models with and without the DGE-IGE covariance can work for GWAS, where either can be chosen *a posteriori* if we observe the inflation or deflation of signals of *p*-values.

### Application to real data

We applied the present model to three datasets — trembling aspen (*Populus tremuloides*), apple (*Malus × domestica*), and grape (*Vitis vinifera*) — to demonstrate its usability. All three datasets involve woody species and provide individual-level data on phenotypes, genotypes, and spatial arrangement. Based on the spatial layout of the study species, we analyzed the influence of nearest neighbors on each trait with field blocks and rows/columns being treated as covariates.

We first analyzed the life-history and defensive traits of aspen genotypes grown in a field garden in Wisconsin, called the WisAsp project [42–44]. Among the 21 trait records collected over two years, the basal area increment at the later growth stage (BAI2024_2015) alone showed a significant negative covariance between DGE and IGE (*χ*^2^ = 4.268 and 5.015, *p* = 0.039 and 0.025 with likelihood ratio tests for 2014 and 2015, respectively: Fig. 3; Table S1). For the late basal area increment, we conducted GP based on 10-fold cross-validation. This GP showed similar performance between models with and without the covariance in terms of *R*^2^ and MAE (Fig. S8), providing consistent evidence with the simulations (Fig. 2c-d and j-k). GWAS of the 21 traits detected a significant IGE SNP for the first bud burst (FDR < 0.05: Table S1), but none for the other traits including defensive chemicals (FDR > 0.05). Besides the plant traits, we analyzed the three quantitative indices of arthropod community composition but found neither significant DGE-IGE covariance nor IGE SNPs (Fig. 3; Table S1). These findings led us to examine intergenotypic competition in other tree species.

**Figure 3.**
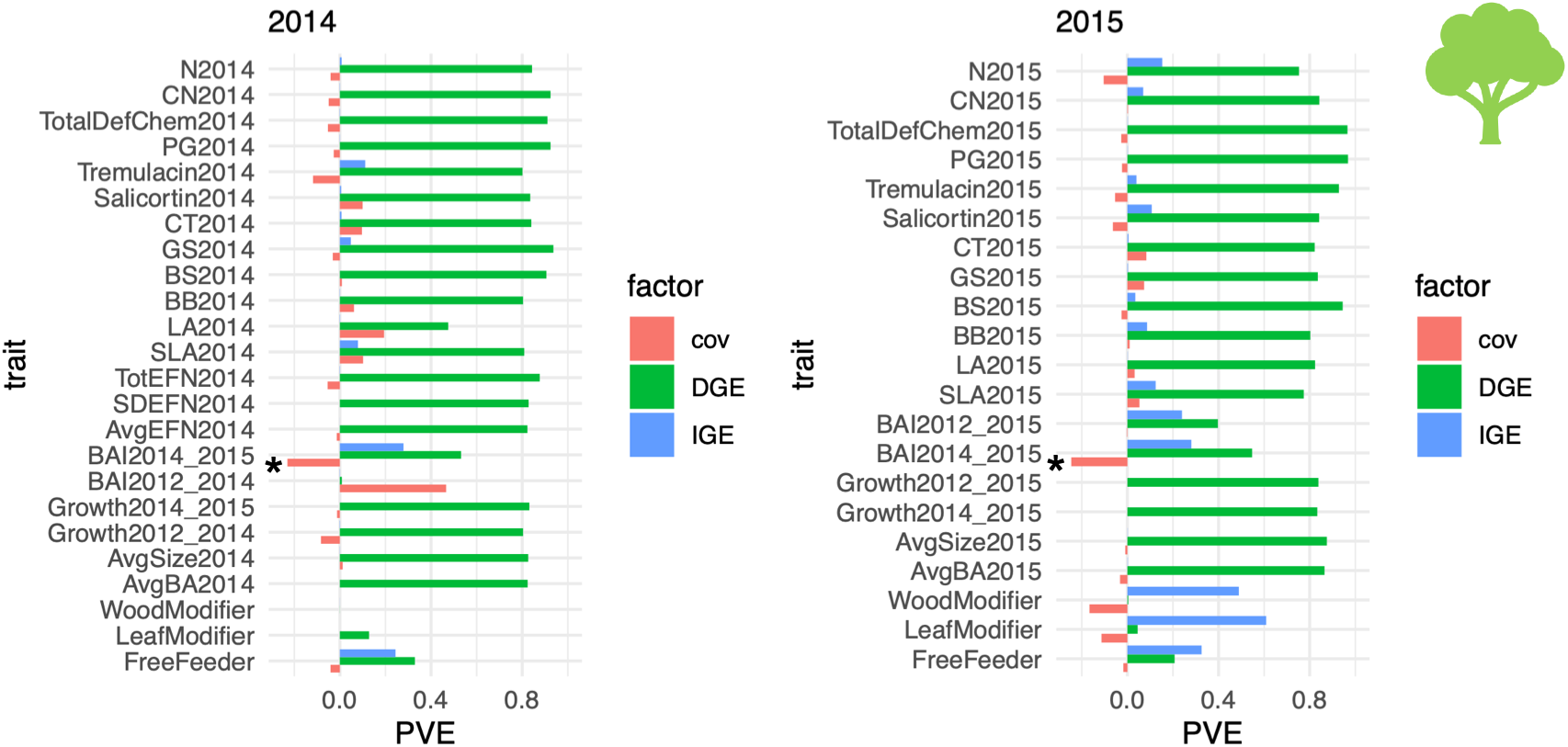
Proportion of phenotypic variation explained (PVE) by direct genetic effects (DGEs), indirect genetic effects (IGEs), and their covariance (cov) in aspen. Blue (upper), green (middle), and red (lower) bars indicate PVE by IGEs, DGEs, and covariance for each trait (see PVE_IGE_, PVE_DGE_, and PVE_cov_ calculations in the main text subsection “Model development”). The left and right panels present results from data collected in 2014 and 2015, respectively. Trait abbreviations: AvgBA, average basal area; AvgSize, average tree volume; GrowthYEAR1_YEAR2, growth of AvgSize from YEAR1 to YEAR2; BAI, basal area increment; AvgEFN, average extra-floral nectary; SDEFN, standard deviation of extra-floral nectary; TotEFN, total extra-floral nectary; SLA, specific leaf area; LA, leaf area; BB, first bud break; BS, terminal bud set; GS, growing season (i.e., BB to BS); CT, condensed tannin; PG, phenolic glycoside; TotalDefChem, total defense chemicals; CN, carbon-nitrogen ratio; and N, nitrogen. Asterisk (^*^) highlights the significant PVE_cov_ by a likelihood ratio test at *p*<0.05. Note that BAI2014-2015 had almost the same phenotypic values between (a) and (b) as it was measured in 2015.

Second, we analyzed apple REFPOP project data collected across multiple years and sites in Europe (Fig. 4; Fig. S6). This dataset included 20,280 individuals of 534 accessions across six European sites for 30 traits [38,45,46]. We quantified PVEs by DGEs, IGEs, and their covariance for these traits (Table S2; Fig. S9) and focused on three traits with fiewer missing records, that is flowering intensity, trunk diameter, and trunk increment (Fig. 4a, c, and e). Flowering intensity was the most largely regulated by DGEs among these three traits (Fig. 4a), whereas the trunk diameter and increment showed relatively large influence of IGEs (Fig. 4c and e). These results suggest that growth traits were more likely to be subjected to IGEs rather than phenological traits. To further examine the intensity of competition among apple trees, we analyzed relationships between PVE by the DGE-IGE covariance (i.e., PVE_cov_) and average tree size (Fig. 4b, d, and f). The flowering intensity showed a positive but non-significant relationship between PVE_cov_ and mean trunk diameter (LMM, slope coef. = 0.0002, *t* = 0.084, *p* = 0.533 by Wald test: Fig. 4b), where we did not detect any significant 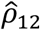 (Table S2). In contrast, the trunk diameter and increment showed significantly negative relationships between PVE_cov_ and mean trunk diameter (slope coef. = -0.004 and -0.007, *t* = - 2.21 and -3.54, *p* = 0.024 and 0.002 for the trunk diameter and increment, respectively: Fig. 4d and f), indicating the intergenotypic competition intensified among bigger trees. Regarding the trunk increment, GWAS also detected two significant IGE SNPs on the seventh chromosome at Belgium site in 2019 (Fig. 4g), which were located on the seventh chromosome at 8743248 and 8744866 base pair positions. Both SNPs had negative *β*_*q*,2_ estimates, indicating the negative influence of allelic mixtures on the trunk increment (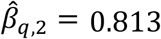: see also Fig. S1 and S2 for the interpretation of ±*β*_*q*,2_). In total, the apple data provide supportive evidence for competitive genetic interactions among neighboring trees.

**Figure 4.**
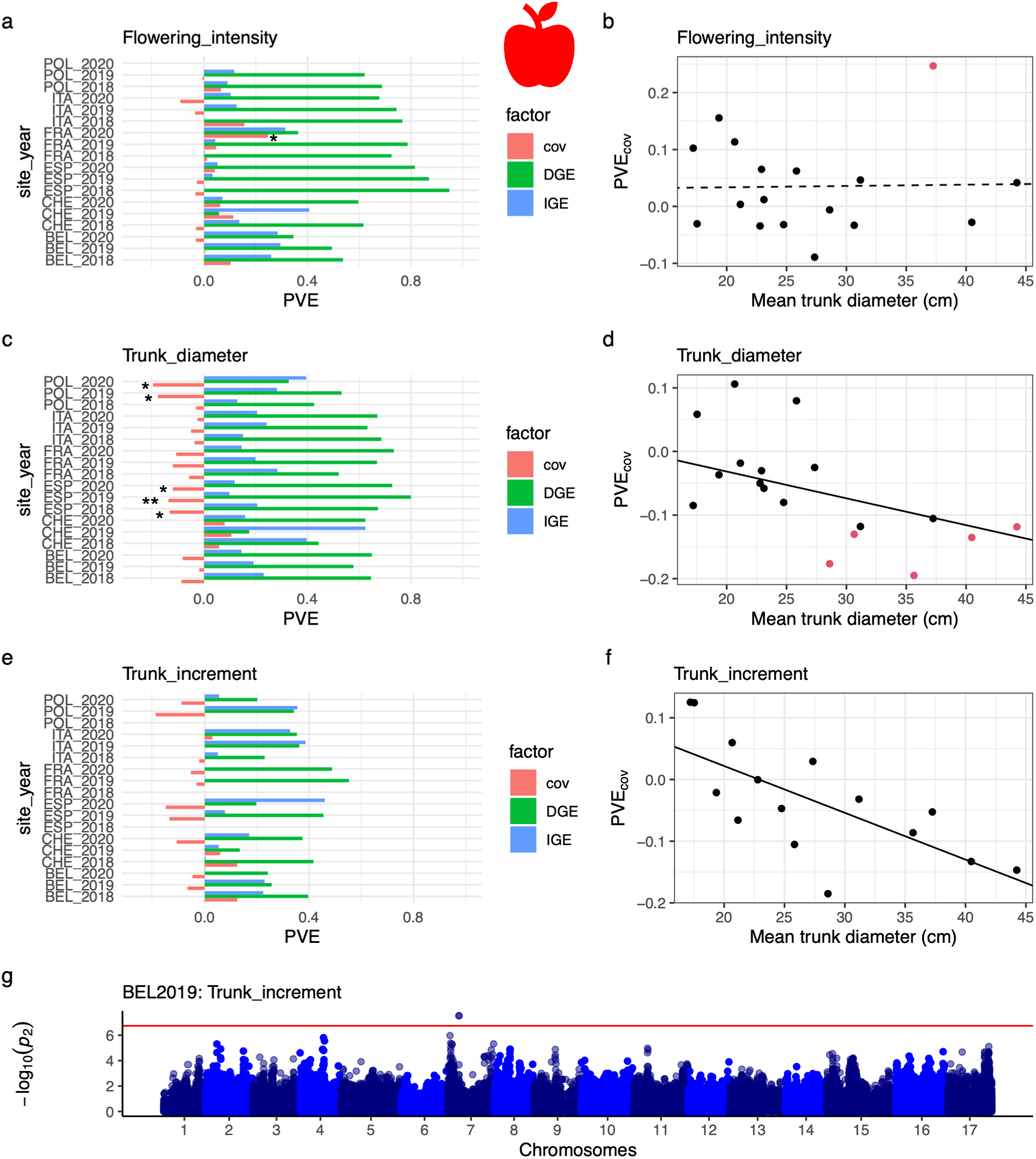
Indirect genetic effects of conspecific neighbors on apple traits across the six sites (Belgium, BEL; France, FRA; Italy, ITA; Poland, POL; Spain, ESP; and Switzerland, CHE) and three years (2018, 2019, and 2020). (a-f) Variation partitioning of the flowering intensity (a-b), trunk diameter (c-d), and trunk increment (e-f). The left panels (a, c and e) show the proportion of phenotypic variation explained (PVE) by direct genetic effects (DGEs), indirect genetic effects (IGEs), and their covariance (cov) following the aspen example (see Fig. 3). The right panels (b, d, and f) display PVE by the covariance against the mean trunk diameter per trial. A single data point corresponds to each field trial. Red circles highlight trials with significant 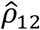. Solid and dashed lines indicate significant and non-significant trends detected by linear mixed models that include the sites and years as random effects. (g) GWAS Manhattan plot of IGEs on trunk increment at the Belgium site in 2019 (BEL2019). The y-axis [-log_10_(*p*_2_)] indicates the association score of *β*_*q*,2_, which is plotted against 17 chromosomes of apple. The horizontal dashed line indicates the genome-wide threshold at FDR = 0.05. Asterisks highlight significant PVE_cov_ by a likelihood ratio test at ^*^ *p*<0.05 or ^**^ *p*<0.01.

Third, we analyzed the grape INNOVINE project data [37] to examine a climbing woody plant as another case. This dataset included 3,642 individuals and 90,007 SNPs for 19 traits at a single site over three years [37]. We separately analyzed each year’s data and found that most traits did not show large influence of neighboring genotypes compared to DGEs (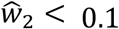 for 17 out of the 19 traits: Fig. 5). Although the number of clusters (nb.clusters) was influenced by IGEs and positive DGE-IGE covariance at the initial stage in 2010 (Fig. 5a), this was not observed in later years (Fig. 5b and c; Table S3). In later years, instead, the pruning weight and citric acid exhibited a significant covariance but not negative covariance with large IGEs. In 2012, we also detected significantly negative covariance for the days-to-fruiting (days.upto.sampling) and shikimic acid, but these traits have little IGE variance compared to DGEs (Fig. 5c; Table S3). These results from climbing grapevines provide limited evidence for competitive interactions among neighboring individuals, highlighting the case in which intergenotypic competitions are difficult to detect.

**Figure 5.**
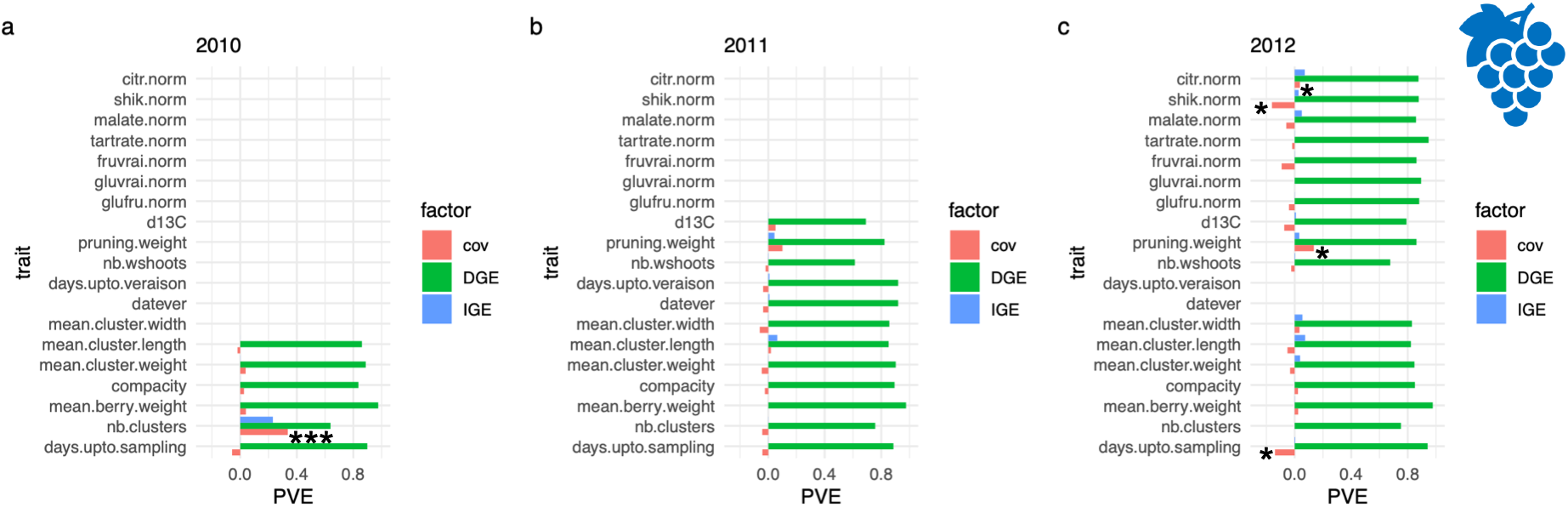
Proportion of phenotypic variation explained (PVE) by direct genetic effects (DGEs), indirect genetic effects (IGEs), and their covariance (cov) in grapes. Blue (upper), green (middle), and red (lower) bars respectively indicate PVE by IGEs, DGEs, and covariance for each trait (see also Fig. 3). Trait abbreviations: “nb.clusters”, the number of clusters; “datever”, date of veraison; “nb.wshoots”, the number of woody shoots, “d13C”, *δ*^13^C; “glufru.norm”, the sum of glucose and fructose; “gluvrai.norm”, glucose; “fruvrai.norm”, fructose; “tartrate.norm”, tartaric acid; “malate.norm”, malic acid; “shik.norm”, shikimic acid; and “citr.norm”, citric acid. Asterisks highlight the significant PVE_cov_ by a likelihood ratio test at ^*^ *p*<0.05 or ^***^ *p*<0.001.

Additionally, we applied the conventional IGE model (Appendix S4) to the three datasets to compare the present and conventional IGE models. Instead of **K**_2_, we defined IGEs by the design matrix, estimated variance component parameters, and tested the significance of the covariance *ρ*_12_ (Table S1-S3). This conventional IGE model detected only a single case of significantly negative *ρ*_12_ in apple, which was detected for the russet frequency at Belgium in 2019 (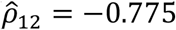, *χ*^2^ = 3.86, *p* = 0.049) yet showed a small contribution by IGEs (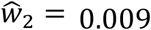: Table S2). We also compared model fitting based on the log-likelihood between the present and conventional models (Eq. 1 and Eq. S4). With the same number of parameters, the present model achieved a larger likelihood than the conventional model on average (Fig. S10). In particular, our model significantly better fitted to the aspen and apple datasets (Wilcoxon signed rank test, *V* = 604 and 25100, *p* = 0.0007 and 1.43e-10 for the aspen and apple, respectively: Fig. S10a and b). These results highlight the cases in which the present model can better account for real data.

## DISCUSSION

Our simulations and applications collectively showed that IGEs and their covariance with DGEs were detectable but did not exceed DGEs. These findings agree with previous evidence that IGEs usually explain <10% of the total phenotypic variation [11,20,47]. Throughout the simulations and applications, we focused on the DGE-IGE covariance *ρ*_12_ as a key parameter underlying the genetic correlations between individual and group performance. Below, we first discuss the methodological features and potential limitations of the present model. Subsequently, we discuss the biological implications of the real data on woody plant species.

### Model performance and limitation

Our simulations showed the performance of the present model in GWAS and GP over our original models. In GWAS, models with or without the covariance *ρ*_12_ showed a similar performance to detect causative IGE SNPs but had a contrasting feature to decrease or increase GIFs depending on the sign of *ρ*_12_ (Fig. S2g; Fig. S5g and n). Practically, we can first try an either of the two models and then consider another if *p*-values are heavily inflated or deflated. This post-hoc screening may help detect candidate genes before experimental validation. Even if we fail to detect significant IGE SNPs, the present model can be used for GP without identifying key SNPs. In GP, the models with the covariance better predicted BLUPs than those without the covariance, although the magnitude of improvement was small when *ρ*_12_ < 0. This was partly because the weighted kernel **K**^′^ sometimes needed to be approximated to a positive definite matrix when *ρ*_12_ < 0, which slightly deviated model estimates and accordingly decreased their BLUP predictability (Fig. S7e). Since SEs became larger when true *ρ*_12_ < 0 (Fig. S5), the negative DGE-IGE covariance seemed more difficult to analyze regardless of the non-positive definiteness. Nonetheless, we were able to detect significantly negative covariance in two out of the three species. This negative covariance co-occurred with smaller IGEs than DGEs (Table S1 and S2), which could be simulated by the biased DGE:IGE ratio and negative covariance (i.e., *w*_1_: *w*_2_ = 0.9: 0.1 with *ρ*_12_ = ™0.6). Together with this simulation scenario, the first application would motivate further analysis of empirical data.

The present and previous IGE models differ in their way of modeling oligogenic and polygenic IGEs. Regarding oligogenic IGEs, the present model aims to test the interactions between DGEs and IGEs at a focal SNP locus by *β*_*q*,2_, which constitutes a form of frequency-dependent selection and thus represents SMA of an allelic mixture [20,31]. Instead, previous GWAS of IGEs tested IGE SNPs without their interaction with DGEs [11,16], which corresponds to *β*_*q*,12_ in the present model and thereby represents GMA. These different choices of fixed effects made it impossible to compare GWAS performance between the present and previous models, but both can be exchanged according to their aim to optimize GMA or SMA. Regarding polygenic IGEs, the largest difference between the present and conventional IGE models lies in the use of the kinship matrix **K** and the design matrix **Z**. The present model splits the kinship matrix into multiple variance-covariance matrices **K**_2_, **K**_12_, and **K**_21_, whereas the conventional IGE models defined polygenic IGEs by splitting **Z** into **Z**_1_ and **Z**_2_ [48]. Since both models possess the same number of (co)variance component parameters, we compared log-likelihood based on real data. Although its performance has yet to be tested more generally, the present model showed a better fit to the aspen and apple data on average (Fig. S10). Thus, our method could provide an alternative tool to analyze IGEs in case the conventional IGE models do not work well.

While the present study focused on the nearest neighboring interactions in continuous two-dimensional space in plants, our model can accommodate two potential extensions. First, it seems plausible for pest spread and volatile-mediated interactions to assume long-range but weaker IGEs from distant neighbors [20,39]. With 0-1 weights included in the design matrix **Z**_2_, previous studies accommodated the spatial distance decay of neighbor genotypic effects [5,49]. Examples of such distance decay include the inverse, exponential, or any other decay functions of Euclidean distance. Although computationally demanding for the models with covariance, the effective range of neighbors can be determined by recursive PVE estimation [30] with **K**_2_ weighted by distance. Second, our model can also be applied to split subpopulations, such as animal herds and potted plants, as analyzed by other IGE studies [11,16]. In this case, our model provides a simplified regression model for a population genetic model of frequency-dependent selection [31] (Appendix S3; Fig. S2 and S3). By applying our model to cage-reared damselflies, previous studies detected negative frequency-dependent selection on female color polymorphism [31]. Practically, this type of grouping can be achieved by setting neighbor spatial scales to infinity and subpopulations as a grouping factor [see 31 and its vignette]. Therefore, the present model based on RAINBOW can be extended to handle split plots and distance decay.

### Genetic architecture of tree competition

To demonstrate the model applicability, we analyzed three woody species, including aspen, apple, and grape. The aspen and apple exhibited negative genetic covariance between DGEs and IGEs during growth. Notably, these negative effects of neighboring genotypes were amplified as trees matured. These results are consistent with previous evidence for significant negative DGE-IGE covariance in *Eucalyptus* and *Pinus* trees [5,25]. In contrast to large trees, grapevines did not exhibit large negative effects of neighboring genotypes on their growth traits. This may be due to their climbing growth form or management practices such as periodical pruning. Regardless of the biological or agricultural reasons, grapes tend to grow vertically and may therefore experience weak competition with horizontal neighbors. Together with the exceptional but plausible case of grapes, the parallel evidence from aspen and apples exemplifies inter-genotypic competition within a plant species.

The present data include phenological and resistance traits, whose patterns might differ from those of growth traits. Regarding phenological traits, flowering time is often explained by DGEs as shown by high heritability within a field site [50,51]. We detected large phenotypic variance explained by DGEs, but limited variance explained by IGEs, for flowering time across the three woody species (e.g., “BS, bud set” in Fig. 3; “Harvesting date” in Fig. S6; and “days.upto.veraison” in Fig. 5). This suggests that phenological traits are unlikely to be explained by IGEs. In contrast to phenological and growth traits, resistance traits or pest damage were expected to show positive DGE-IGE covariance by preventing pest spread to neighbors [[5]; [20]; see also Appendix S6]; however, we could not find significantly positive DGE-IGE covariance regarding secondary metabolites and pest damage. Given that our simulations better detected positive than negative covariance, further applications to resistance traits are needed to test whether positive DGE-IGE covariance can be detected from real data.

Together with the analysis of (co)variance components, GWAS detected two site-specific but significant IGE SNPs with negative 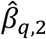 for the apple trunk increment. The negative sign of these SNPs suggests that neighbors with dissimilar alleles reduce the trunk increment of focal individuals, providing evidence for intergenotypic competition at allelic levels. These two significant SNPs exhibited a complete linkage disequilibrium (*R*^2^ = 1) and were closely located on the top-to-center positions of seventh chromosome. For this chromosomal region, previous publications documented QTLs underlying DGEs on fruit size [52] and basal trunk increment [53]. More specifically, our significant SNPs range between 8743248 and 8744866 bp positions on the seventh chromosome, which include a locus encoding a protein kinase superfamily protein (MD07G1084900). This locus is homologous to AT5G56790 in *A. thaliana*, but its biological function remains unknown. The nearest loci of MD07G1084900 include the homologs of MYB33 (MD07G1085000; homolog of AT5G06100) and SUCROSE PHOSPHATE SYNTHASE 1F (MD07G1084800; homolog of AT5G20280).

Although these two genes AT5G06100 and AT5G20280 are involved in the vegetative phase change [54] and leaf CO_2_ assimilation [55] in *A. thaliana*, respectively, their functions have yet to be validated in apples. To this end, our study uncovered a QTL and candidate genes underlying intraspecific competition.

Our method is widely applicable to a broad range of plant GWAS/GS data as far as genotypes are randomly cultivated in two-dimensional space. Such a randomized block design keeps genotypes and groups uncorrelated, thereby allowing us to treat blocks as non-genetic fixed effects [56]. Furthermore, plant data enable us to utilize the spatial positions of field columns and rows as non-genetic covariates to separate neighbor genotypic effects from spatial trends [39,57]. While the present data include individual trees as a unit of genotype [37,38,42], our method can treat multiple individuals of a single genotype, such as a single plot of field crops, as a unit of genotypes [39]. Even when genotypes are spatially split into pots or plots without the continuous positions of rows and columns, shared environments can be analyzed as non-genetic random effects called indirect environmental effects [3,11,16]. This holds if kins constitute social groups, where fixed effects cannot separate IGEs and groups [58,59]. To adequately consider spatial heterogeneity, we would need to include additional fixed or random effects depending on the layout of genotypes and groups.

## Conclusion

The present study enabled GP and GWAS of IGEs by modeling the oligogenic and polygenic effects of neighboring genotypes on a complex phenotype. This method can be kept available as an open-source R package by leveraging RAINBOWR and rNeighborGWAS packages [30,34], which offer an alternative implementation to a licensed ASReml-R package [60], mixed-model solver named “sommer” [61], and a Python-based LIMIX software [62]. While IGEs have been investigated as a theory of social evolution [2,63], empirical applications have mainly been conducted in domesticated animals and plants [1,4,49,64]. Consequently, the genomics of IGEs promises to contribute to both fundamental and applied issues, including multi-level selection and its agricultural application. Recent studies have highlighted a growing demand for an interdisciplinary approach to IGEs [3,12,23]. With the increasing availability of genome-wide polymorphism data, our method may enhance interdisciplinary practice by facilitating GWAS/GP of IGEs across diverse biological systems.

## METHODS

### GWAS/GP implementation

We used the RAINBOWR (>v0.1.42) [34] and rNeighborGWAS (>v1.2.4) [30] packages to implement the multi-kernel mixed model (Eq. 1). Variance and covariance component parameters were estimated using the restricted maximum likelihood (REML) method without the fixed effects *β*_*q*,1_, *β*_*q*,2_, and *β*_*q*,12_. The statistical significance was determined using likelihood ratio tests by dropping *ρ*_12_ out of the full model. The mixed models with or without *ρ*_12_ were solved using the EM3.cpp or EM3.cov functions of the RAINBOWR package [34], respectively. When the kernels did not become semi-positive definite, they were approximated by the nearest positive definite matrix using the nearPD function in R (which was enabled with the “forceApproxK=TRUE” option in the EM3.cov function). In addition to the statistical significance, we calculated SEs of (co)variance components from Fisher’s information matrix (Appendix S5), which was implemented as the calcParamSE.cov function in the RAINBOWR package. For GWAS, each SNP locus was subjected to a likelihood ratio test using an eigenvalue decomposition and diagonalization method for the weighted kernel **K**^′^, which was implemented in the lmm.diago function of the gaston package [65]. This GWAS procedure followed the nei_imm function of the rNeighborGWAS package [30]. All source codes and input data are deposited on the GitHub and Zenodo repositories (see “Data accessibility” section), which includes the vignette and subroutine functions for GWAS/GP implementations. An easy R package to implement Eq. 1 is also available at GitHub and Zenodo repositories (see “Data accessibility” section).

### Simulation

We conducted simulations to examine the performance of the present model, as follows: (i) simulating genomes and phenotypes, (ii) assessing BLUP predictability and causal variant detection, and (iii) comparing these abilities between the models with and without the covariance *ρ*_12_. We utilized previously simulated genotype data [31]. Specifically, by simulating spatial and temporally varying selection over 2,000 generations among ten subpopulations, Sato et al. [31] generated 30 independent datasets, each comprising 2,000 diploid individuals with 50 kb nucleotide sequences across three chromosomes (see also Appendix S4 of Sato et al. [31]). This simulation was performed using SLiM version 3 [66] and its outputs were deposited as variant call format (.vcf) on Dryad (https://doi.org/10.5061/dryad.zs7h44jdv). Depending on the iterations, each genotype dataset contained approximately ten causal DGE SNPs out of ca. 2,500 SNPs (see Figure S4b and c of Sato et al. [31]). Out of wider parameter range and combinations than described in the present study, our previous study [30] identified conditions under which the model performance differentiated. Our base parameters therefore followed those of our previous study [30].

### Variance component and genomic prediction

To test the model’s capability in the inference and prediction of polygenic IGEs, we simulated phenotypes under two scenarios, i.e., negative and positive covariance between polygenic DGEs and IGEs with their even or biased contributions. Specifically, we assumed the true parameters of *ρ*_12_ = 0, ±0.3, or ±0.6 with PVE_u_ = 0.6; *w*_1_: *w*_2_ = 0.5: 0.5 or 0.9:0.1; and 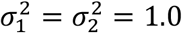. With these true parameters, we randomly assigned 1,600 individuals to a 40 × 40 lattice space. We then simulated a phenotype subjected to DGEs and IGEs from the nearest neighbors along off-diagonals (i.e., 1 Euclidean distance from a focal individual). To test the applicability to genomic prediction, we divided the simulated data into training and test datasets. One-tenth of the individuals were randomly left out of the simulated data as a test dataset, whereas the rest was fitted by the present model as a training dataset. The training model incorporated the linear and quadratic terms of spatial rows and columns as non-genetic covariates. To determine the trait predictability using the test dataset, we calculated the coefficient of determination (*R*^2^) and MAEs between the estimated and simulated BLUPs. This line of simulations was repeated for the 30 genotype datasets as described above.

### Genome-wide association study

We also tested the model performance to detect causative IGE SNPs in GWAS. Our procedure followed Sato et al. [31] while simulating the negative or positive covariance between DGEs and IGEs (*ρ*_12_ = ±0.6) under even or biased DGE:IGE ratio (*w*_1_: *w*_2_ = 0.5:0.5 or 0.9:0.1). The other parameters were the same as those described above for GP. We also assumed 20 causative IGE SNPs, of which ten had a positive coefficient as *β*_*q*,2_ = 0.1 and the remaining ten had a negative coefficient as *β*_*q*,2_ = ™0.1. To make DGE and IGE correlated, we further assumed that the causative IGE SNPs overlapped with DGE SNPs (i.e., *β*_*q*,1_ = *β*_*q*,2_ = ±0.1). This assumption was not so critical, as our conclusion did not change even when causative SNPs were independent between DGEs and IGEs (Fig. S11). In addition to non-zero *β*_*q*,1_ and *β*_*q*,2_, half of the causative IGE SNPs were assumed to have non-zero *β*_*q*,12_ as *β*_*q*,12_ = ±0.1. With or without *ρ*_12_, the present model Eq. 1 was fitted to simulated phenotype **y** to estimate the weighted kinship matrix **K**^′^. Based on the RAINBOW algorithm [34], we applied eigenvalue decomposition to **K**^′^ to correct for population structure in a background mixed model. To distinguish between DGE and IGE SNPs, we performed GWAS of *β*_*q*,2_ using a likelihood ratio test by dropping *β*_*q*,2_ out of the full model including both DGEs and IGEs from a focal SNP (i.e., *β*_*q*,1_ and *β*_*q*,2_). After model fitting, GWAS performance was evaluated based on the false positive and negative rate to detect causative IGE SNPs. The balance between false and true positive rates was calculated using the receiver-operating characteristic curve (ROC) and its area under the curve (AUC). These ROC curves were also used to determine the power to capture true positives at the stringent false discovery rate (FDR) = 0.05. We utilized the pROC package implemented in R [67] for the ROC and AUC analysis. To examine the risk of false positive and negative signals, we calculated genomic inflation factors [40,41] as 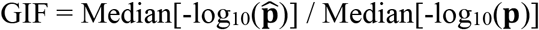 where 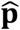 and **p** indicate the vector of observed and randomly expected *p*-values, respectively. Accordingly, GIFs of >1.0 and <1.0 indicate the inflation and deflation of *p*-values, respectively.

### Application

#### Aspen

For WisAsp data, we curated phenotype and genotype data from Barker et al. [42] and [43], respectively. These data were deposited at Dryad (https://doi.org/10.5061/dryad.fr045hv and https://doi.org/10.5061/dryad.st463). The annotation of genotype-phenotype link was obtained from Riehl et al. [44]. We set the cut-off value of minor allele frequency (MAF) at 0.05. In total, we analyzed data from 813 individuals in 2014, comprising 169,150 SNPs, and data from 826 individuals in 2015, comprising 169,111 SNPs. In this SNP matrix, 73% of the loci were homozygous while the remaining 27% were heterozygous.

Using the curated aspen data, we performed variance component analysis and GWAS of IGEs. The influential area of neighbors was assumed for the nearest neighbors along the off-diagonal direction, i.e., 1 Euclidean distance from a focal individual. We included experimental blocks as non-genetic factorial covariates in all the mixed models. To consider non-genetic spatial covariates, we also incorporated the field rows and columns whose linear and quadratic terms were expected to represent environmental gradients and edge effects, respectively. These analyses were separately performed for the years 2014 and 2015. For each trait per year, we tested the statistical significance of the covariance parameter *ρ*_12_ over the model with *ρ*_12_ = 0 based on the likelihood ratio test. According to the estimated variance components, we then conducted GWAS of *β*_*q*,2_ based on the weighted kinship matrix **K**^′^. Unless otherwise stated, we determined the significance threshold by FDR for GWAS. We used the CalcThreshold function implemented in the RAINBOWR package [34] to calculate Benjamini-Hochberg’s FDR [68] for GWAS. We also calculated GIFs for all traits but found no signals of *p*-value inflation at >1.2 except for wood modifiers in 2015 (which were heavily zero-inflated and exceptionally showed a positive log-likelihood: see Table S1).

#### Apple

We curated the apple REFPOP data available via the Data INRAE (https://doi.org/10.15454/VARJYJ) and Zenodo (https://zenodo.org/records/14191209) repositories [38,46]. The phenotype data deposited by Jung et al. [38] included 30 traits for 20,280 individuals of 534 accessions across six European sites (Belgium, France, Italy, Poland, Spain, and Switzerland). Ten of these traits were measured at one location, while the remaining 20 traits were available from at least two locations (cf. Fig. 1 of Jung et al. [45]). Genotype information of the 534 accessions is available as imputed SNP data between 20K and 480K array [45], yielding 281,140 SNPs at MAF > 0.05. In the SNP matrix, 66% of the loci were homozygous while the remaining 34% were heterozygous. The imputed genotype data are deposited by Jung et al. [46] on Zenodo as described above. When screening candidate genes after GWAS, we referred to the whole genome assembly and annotation of *Malus x domestica* GDDH13 v1.1 [69] through the Apple Genome and Epigenome project website (https://iris.angers.inra.fr/gddh13/; Last accessed on 18-Feb-2026).

Using the curated apple data, we conducted variance component analysis and GWAS per year and site. In this dataset, individuals were assigned to rows and positions. Depending on the site, planting distance within and between tree rows ranged 0.9-1.3 m and 3.2-3.6 m, respectively [45]. Therefore, we considered the nearest neighboring interactions along the position within the tree rows. We also included the linear and quadratic terms of rows and positions as non-genetic spatial covariates in all the mixed models. For each year and site, we then performed a likelihood ratio test for *ρ*_12_ and GWAS of *β*_*q*,2_ as described for the aspen data above. To test whether PVE by DGE-IGE covariance (PVE_cov_) decreased as the mean trunk diameter increased, we used linear mixed models in which the study years and sites were considered random effects. We also calculated GIFs for all traits per sites and years but found no signals of *p*-value inflation at >1.2 (Table S2).

#### Grape

We downloaded the set of grape phenotype and genotype data from the Data INRAE repository (https://doi.org/10.15454/8DHKGL), which was archived by Flutre et al. [37]. These data comprise 3,642 individuals of 270 accessions with 90,007 SNPs at MAF > 0.05. Of these SNP loci, 79% were homozygous while the remaining 21% were heterozygous. Nineteen traits were available, some of which exhibited numerous missing data points (Table S3) (see also Supplementary Fig. 1 of Flutre et al. [37]). During data curation, we found an outlier for citric acid content (citr.norm), which exhibited a 100-times larger phenotypic value compared to the other, and thus excluded this data point from the subsequent analysis.

The retrieved grape data contained trait records over three years (2010, 2011, and 2012); therefore, we separately analyzed each trait per year for variance component analysis and GWAS. In this dataset, individuals were assigned to ranks and locations in the field, where the ranks were separated 2.5 m each other while the locations were 1 m apart between individuals within a rank [37]. Therefore, we assumed he nearest neighboring interactions along locations with ranks considered a grouping factor. We also included the experimental blocks as non-genetic factorial covariates while the linear and quadratic term of ranks and locations as non-genetic spatial covariates. Then we performed likelihood ratio test for *ρ*_12_ and GWAS of *β*_*q*,2_ as described for the aspen and apple data above. We also calculated GIFs for all traits per year but found no signals of *p*-value inflation at >1.2 (Table S3).

## Supplementary materials

## Appendices Appendix S1. Original model of Neighbor GWAS

In the main text, we modified the original Neighbor GWAS model to implement a flexible mixed model including polygenic and oligogenic IGEs. Here, we present the original model [30] to highlight how to model locus-wise IGEs. Following the main text (Fig. 1), we modeled the IGEs of neighboring genotypes at the *q*-th SNP locus on focal *k*_*i*_-th individual phenotype 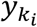 at the local space *k* as follows:

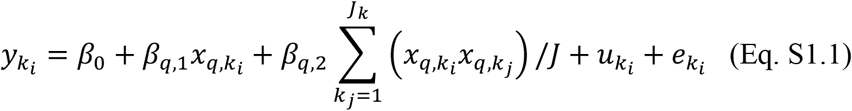

where *x*_*q,i*_ and *x*_*q,j*_ represent the genotype of focal *i*-th and neighbor *j*-th individuals in a local space *k* (where *J*_*k*_ is the total number of neighboring individuals). For simplicity, let us assume inbred lines with two homozygotes are encoded as aa = -1 and AA = +1 Then, the fixed effect *β*_*q*,2_ indicates the influence of mean allelic (di)similarity [i.e., (-1)×(+1) = (+1)×(- 1) = -1 or (+1)×(+1) = (-1)×(-1) = +1] between focal and all neighboring individuals on 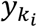. This is analogous to the Ising model of ferromagnetics [35], in which *β*_*q*,2_ represents physical interactions among the south/north dipole of neighboring magnets while *β*_*q*,1_ is an external energy. By focusing on the fixed effects of Eq. S1.1, Figure S1 exemplifies spatial patterns shaped by neighbor genetic interactions over time, in which negative and positive *β*_*q*,2_ favors spatial mixing and separation of two genotypes in a neighborhood (Fig. S1a and b, respectively). In real data, we aim to estimate *β*_*q*,1_ and *β*_*q*,2_ by regressing observed 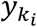 on 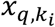.

In addition to the fixed effects *β*_*q*,1_ and 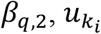 denotes random effects that represent polygenic self and neighbor genotypic effects, respectively. Using vectors and matrices, Eq. S1 can also be expressed as a mixed model as follows:

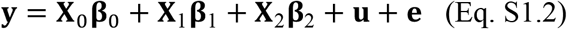

In the original model [30], **u** consists of additive polygenic effects from self and neighbor genotypes as 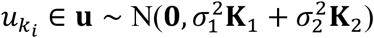. The residual *e*_*i*_ is defined as *e*_*i*_ ∈ **e** and 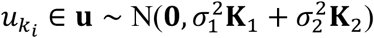. **Z** denotes a design matrix and can be an identity matrix **I** when all genotypes are different. The *N* × *N* variance-covariance matrices **K**_1_ and **K**_2_ are the same as those defined in the main text (see “Model development”).

When the two random effects **u**_1_ and **u**_2_ are additive and independent, the total phenotypic variance can be decomposed as 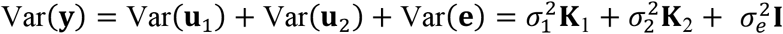. The total genetic and environmental variance can then be partitioned as 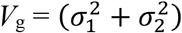 and 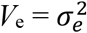. The proportion of phenotypic variation explained by additive DGEs and IGEs (PVE_1+2_) is then given as PVE_1+2_ = *V*_g_ / (*V*_g_ + *V*_e_). Furthermore, the net contribution of IGEs to the phenotypic variation should be calculated as 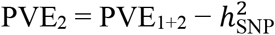, where 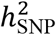 denotes the conventional SNP heritability. Although this conservative metric was needed due to a strong element-wise correlation between **K**_1_ and **K**_2_ [30], this issue would be addressed by allowing the covariance parameter *ρ*_12_ = (™∞, ∞) to decrease or increase the total genetic variance in the present study (see Eq. 2).

## Appendix S2 Interaction term between SNP-wise self and neighbor effects

The original model Eq. S1.1 implicitly assumes symmetric effects between focal and neighboring individuals at the locus *q* by 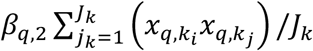. This assumption can be relaxed by introducing an interaction term between DGE and IGE as follows:

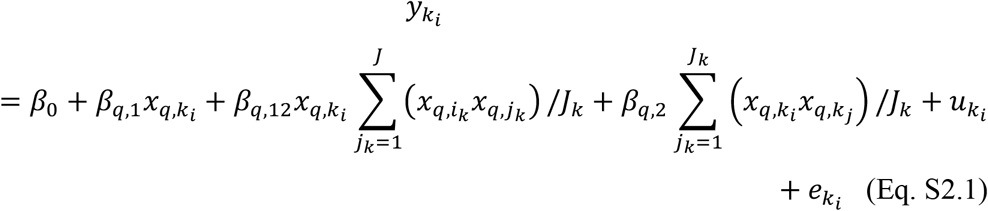

When genotype values are encoded as {AA, Aa, aa} = {+1, 0, -1}, squared-genotype values 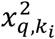 become +1 or 0. This leads to 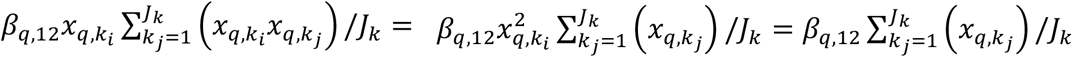, thereby simplifying Eq. S2.1 as follows:

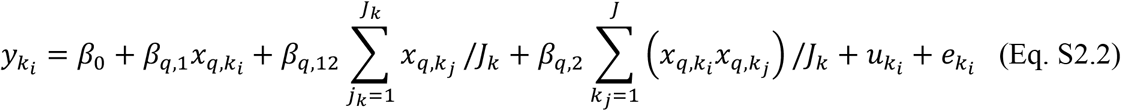

This Eq. S2.2 indicates that in the context of the Ising model, the statistical interaction term *β*_*q*,12_ can be exchanged as a main effect [70]. For clarity, we redefine *β*_*q*,1_ = *β*_*q*,s_; *β*_*q*,12_ = *β*_*q*,n_; and *β*_*q*,2_ = *β*_*q*,s×n_, which yields:

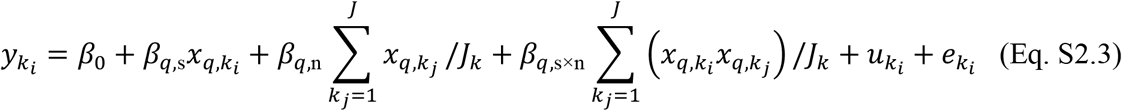

Similarly, Eq. S2.3 can be extended as

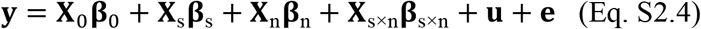

where the three random effects **u**_s_, **u**_n_, and **u**_s×n_ can be defined as 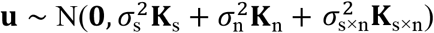. To correct for the population structure due to asymmetric neighbor effects, Eq S2.4 introduced the third random effect into the original model. Similar to **K**_s_ = **K**_1_ and **K**_s×n_ = **K**_2_, we can define **K**_n_ as the *N* × *N* variance-covariance matrix that represents polygenic self-by-neighbor genotypic effects on 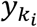. This self-by-neighbor matrix **K**_n_ is also given by the cross-product as 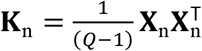. The elements of the *N* individuals × *Q* SNPs matrix **X**_n_ consist of covariates for the self-by-neighbor genotypic effects as 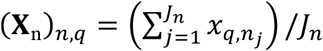. After estimating the three variance component parameters 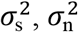, and 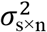, we can obtain a weighted kernel as 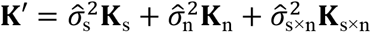. Similar to GWAS based on Eq. 1 (see Methods), this weighted kernel matrix represents the total population structure due to polygenic self, neighbor, and self-by-neighbor effects. GWAS can be performed using an eigenvalue decomposition of the weighted kernel **K**^′^ and diagonalization method implemented in the lmm.diago function of the gaston package [65].

## Appendix S3 Theoretical expectation of mean phenotypic values

In terms of evolutionary biology, the Neighbor GWAS model offers a straightforward regression method for population genetic models of frequency-dependent selection [31,70], also known as pairwise interaction models [71]. Specifically, the sign of *β*_*q*,2_ represents positive or negative frequency-dependent selection when 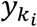 is a fitness-related phenotype [31]. In other words, a sort of Eq. S1.1 and S2.1 pose a selection gradient analysis of frequency-dependent selection to estimate *β*_*q*,2_ by regressing phenotypes on genotypes and their spatial similarity [31].

By focusing on allele frequency within a local space *k*, we can clarify how the three locus-wise coefficients *β*_*q*,s_, *β*_*q*,n_, and *β*_*q*,s×n_ affect the focal phenotypic value 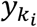 [31]. Given that all the three datasets analyzed in this study included a non-negligible fraction of heterozygotes (see Methods), we assumed additive effects and encoded genotype values as {AA, Aa, aa} = {+1, 0, -1}. This holds the case 2 in Appendix S2 of Sato et al. [31]. According to Sato et al. [31], let us assume random mating and interactions among AA, Aa, and aa genotypes in the local space *k* where *f*_*q*,A_ denotes the frequency of A alleles at *q*-th SNP locus. The phenotypic values for AA, Aa, and aa genotypes are then given as follows:

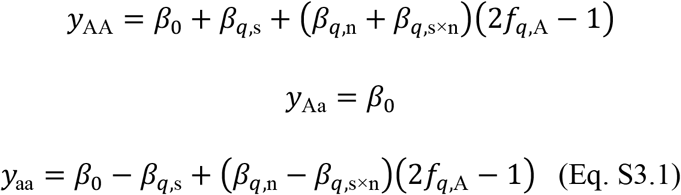

Then, the marginal phenotypic values at allele levels are given by their weighted mean as *y*_A_ = *β*_0_ + *f*_*q*,A_(*β*_*q*,n_ + *β*_*q*,s×n_)(2*f*_*q*,A_ ™ 1) + *β*_*q*,s_*f*_*q*,A_ and *y*_a_ = *β*_0_ + (1 ™ *f*_*q*,A_)(*β*_*q*,n_ ™ *β*_*q*,s×n_)(2*f*_*q*,A_ ™ 1) ™ *β*_*q*,s_(1 ™ *f*_*q*,A_) (see also Appendix S2 of Sato et al. [31]). Furthermore, the group mean of 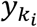 is given by the weighted mean between the two genotypes as 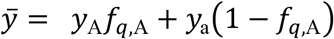. According to Eq. S3.1, Figure S2 presents numerical examples of *y*_A_, *y*_a_ and 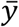 in response to *f*_*q*,A_. When *β*_*q*,n_ < 0, the group mean 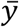 decreases as increasing *f*_*q*,A_ (Fig. S2c and f). Contrarily, 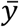 increases as increasing *f*_*q*,A_ when *β*_*q*,n_ > 0 (Fig. S2b and e). These two cases indicate that the sign of *β*_*q*,n_ determines a positive or negative slope of the group mean along *f*_*q*,A_. Regarding the other coefficients, *β*_*q*,s_ modulates the intercept while *β*_*q*,s×n_ determines relative difference (i.e., frequency-dependent selection) between *y*_A_ and *y*_a_ and resulting shape of 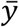 (i.e., convex or concave: Fig. S2a and d). These examples showcase the way in which the group mean 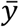 decreases or increases in response to frequency-dependent selection on 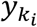.

The additive case converts genotype values as {AA, Aa, aa} = {+1, 0, -1} and thereby makes interactions involving heterozygotes become zero, i.e., *x*_*q*,AA_*x*_*q*,Aa_ = *x*_*q*,Aa_*x*_*q*,Aa_ = *x*_*q*,aa_*x*_*q*,Aa_ = 0. To relax this simplification, we also examined the half-similarity case with *x*_*q*,AA_*x*_*q*,Aa_ = 0.5 and *x*_*q*,aa_*x*_*q*,Aa_ = ™0.5. The phenotypic values for this case are then given as follows:

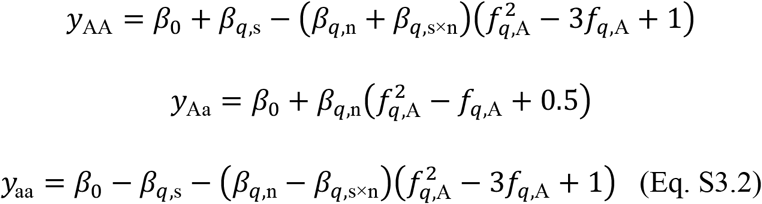

For Eq. S3.2, the marginal phenotypic values at the allele levels are given by their weighted mean as *y*_A_ = *f*_*q*,A_*y*_AA_ + (1 ™ *f*_*q*,A_)*y*_Aa_ and *y*_a_ = *f*_*q*,A_*y*_Aa_ + (1 ™ *f*_*q*,A_)*y*_aa_. The fitness difference between A and a alleles is 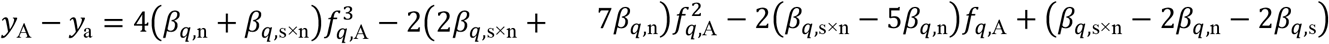. Solving *y*_A_ ™ *y*_a_ = 0 with respect to *f*_*q*,A_ yields three potential equilibria at the maximum if Cardano’s formula is applied to the cubic equation. Furthermore, the weighted mean 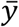 is defined by 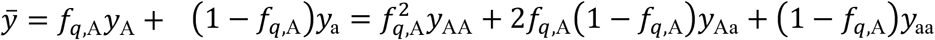, which yields a quartic function of *f*_*q*,A_ as 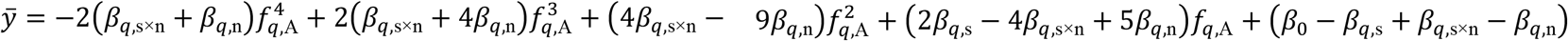. Figure S3 presents numerical examples for this quartic function of *f*_*q*,A_, showing that the sign of *β*_*q*,s×n_ is a key to the concave or convex shape of 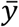 while *β*_*q*,s×n_ gives a slope along *f*_*q*,A_. Therefore, the present study adopts a simpler additive case than the half-similarity.

In the main text, the present model Eq. 1 extends *β*_*q*,s_, *β*_*q*,n_, and *β*_*q*,s×n_ to accommodate their polygenic effects. As shown in Figure S2, the sign of *β*_*q*,n_ (= *β*_*q*,12_) determines whether the group mean decreases or increases when the AA genotypes are selected to become rare or abundant in the group *k*. According to the interpretation of *β*_*q*,n_ (= *β*_*q*,12_), the covariance *ρ*_12_ determines the negative or positive response of self-polygenic effects 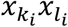 to the neighboring allele frequency 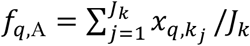 (see also the main text Eq. 1). Taken together, the covariance *ρ*_12_ can be interpreted as a key parameter that represents negatively or positively correlated responses between polygenic DGE and IGE. This interpretation aligns with the DGE-IGE covariance in the ordinal IGE model (see Appendix S4 below).

## Appendix S4 Relationship with other IGE models

Regarding random effects, conventional IGE models designate neighboring individuals by a design matrix or incidence matrix **Z** [5,11,48]. These conventional models can also be implemented using the RAINBOWR package as it allows a user-defined **Z**. Here, let the two design matrices be **Z**_1_ and **Z**_2_ for polygenic DGEs and IGEs, respectively. The ordinal model is then given as follows.

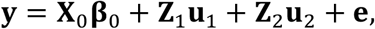

where

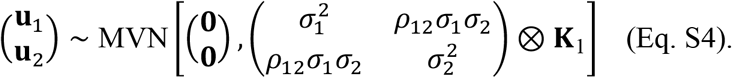

This ordinal model defines polygenic DGE and IGE by random effects **u**_1_ and **u**_2_ without fixed effects for each SNP. In the random effects, the kinship matrix **K**_1_ is shared among the variance and covariance parameters 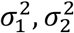, and *ρ*_12_ as denoted by the Kronecker product ⊗. In this context, the sign of the covariance *ρ*_12_ determines antagonistic (negative) or synergistic (positive) responses of focal and neighboring individuals to selection [48].

Regarding fixed effects, previous studies implemented SNP-wise fixed effects from focal and neighboring individuals with Eq. S4 set as a background null model [e.g., 11,16]. To implement GWAS of IGEs, these studies modeled SNP-marker effects from neighbor genotypic values [11] or mean genotypic values among group mates [16], which corresponded to what *β*_12_ represented in our study. Our GWAS can be exchanged into other IGE GWASs by testing *β*_12_ instead of *β*_2_, although comparing the GWAS performance between ours and previous GWAS remains difficult due to their different choice for *β*_12_ or *β*_2_.

## Appendix S5 Standard errors of (co)variance components

To corroborate the results of the simulations and applications, we calculated the standard errors (SEs) of the variance component parameters. According to Searle et al. [72], the asymptotic standard errors of variance component parameters can be calculated from the diagonal elements of the inverse Fisher’s information matrix 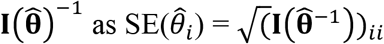, where 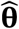 means the vector of estimated variance component parameters 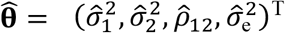. The elements of Fisher’s information matrix are defined by the expected values of the negative Hessian matrix, which consists of the second derivative of a negative log-likelihood function [72,73]. When **Z** = **I**, Fisher’s information matrix is given as follows (see Eq. 94 and its derivation at page 253 of Searle et al. [72]):

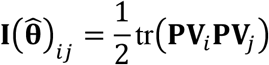

where tr(…) denotes the trace of a matrix. The variance-covariance matrix **V** denotes the total variance and **V**_*i*_ refers to its first derivative with respect to the *i*-th parameter *θ*_*i*_, i.e., **V**_*i*_ = ∂**V**/ ∂*θ*_*i*_. The projection matrix **P** is idempotent (i.e., **PP** = **P**) and is defined by **P** = **V**^-1^ ™ **V**^-1^**X**(**V**^T^**V**^-1^**X**)^-1^**X**^T^**V**^-1^) with the matrix **X** indicating the intercept and non-genetic covariates.

Then, let us consider the case of Eq. 1 without SNP-marker effects as **y** = **X**_0_**β**_0_ + **u**_1_ + **u**_2_ + **e**. The total variance for Eq. 1 is designated as 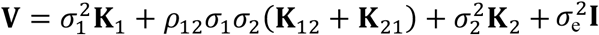. Partially differentiating **V** with respect to 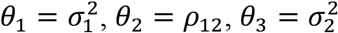, and 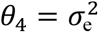, yields four derivatives as follows.

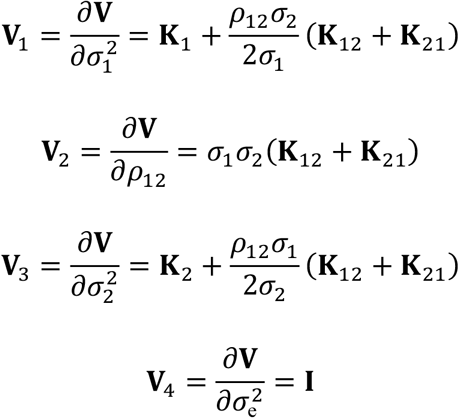

Substituting 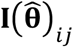 by **V** to **V** gives 4 × 4 Fisher’s information matrix, from which we can calculate SEs of the *i*-th parameter following 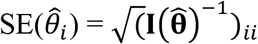 . Although these asymptotic SEs are not equivalent to unbiased SEs corrected by group size [74], they tend to be larger than true errors when DGE and IGE are strongly correlated [see also 59 for simulations]. Therefore, we performed a likelihood ratio test in case the asymptotic SEs underestimated significance levels by increasing confidence intervals. As described in the main text, these asymptotic SEs exhibited consistent variation with the simulated estimates of (co)variance components (Fig. S6). When 1.96×SEs can be considered 95% confidence intervals, these intervals showed consistent 5% significance levels with those determined by likelihood ratio tests for the real data (Table S1-S3).

## Appendix S6. Revisiting field herbivory on *Arabidopsis thaliana*

As discussed in the main text, disease phenotypes are reported to have positive covariance between DGEs and IGEs [5]. Previously, we applied our original model and detected the significant contribution of neighboring genotypes to herbivore damage and community composition on field-grown 199 *A. thaliana* accessions [20]; however, our original model did not consider the covariance between the self and neighbor genotypic effects. Here, we revisited our published data using the present model. The original data include four phenotypes for nearly 6,400 individual plants (∼ 199 accessions × 8 randomized blocks × 2 sites × 2 years), such as the herbivore damage, the individual number of external and internal feeders, and the total number of insect species in Switzerland and Japan [75]. Except for the DGE-IGE covariance, we repeated the same analysis of the first nearest neighbors as in our previous work. As a result, this reanalysis detected a significantly positive DGE-IGE covariance for the herbivore damage in Switzerland (*ρ*_12_ ± SE = 0.434 ± 0.148; likelihood ratio test, 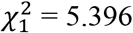, *p* = 0.020) while the other phenotypes did not manifest any significant covariance (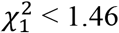, *p* > 0.22). The herbivore damage evaluated by the number of leaf holes in Switzerland yielded a fraction of total genetic variance 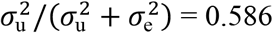. This fraction exceeded the previously reported values of 0.513 by our original model as well as the SNP heritability of 0.453 (see Figure 2e of Sato et al. [20]). These additional findings provide a case in which the positive DGE-IGE covariance amplifies heritable variation, paving the way for further applications to pest damage and defensive traits.

## Supplementary Tables

Table S1. Variance component estimates and significant IGE SNPs in aspen. The effective number of individuals with trait records (*N*), DGE variance 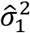, IGE variance 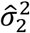, DGE-IGE covariance 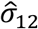, environmental variance 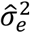, log-likelihood (LL), likelihood ratio and its p-values over a model without the covariance (*χ*^2^ and *p*), the number of significant IGE SNPs (#sig. SNPs), and genomic inflation factor (GIF) are shown for 2014 and 2015. The results of the conventional IGE model (Eq. S4) are shown on the right, which include LL, IGE weight 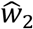, DGE-IGE covariance 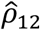, likelihood ratio *χ*^2^ and *p*-values over the model without *ρ*_12_. The trait abbreviations are available in the caption of Figure 3. NA in the standard error (SE) means not applicable due to too small or large estimates.

Table S2. Variance component estimates and significant IGE SNPs in apple. The effective number of individuals with trait records (*N*), DGE variance 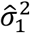, IGE variance 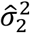, DGE-IGE covariance 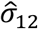, environmental variance 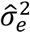, log-likelihood (LL), likelihood ratio and its p-values over a model without the covariance (*χ*^2^ and *p*), the number of significant IGE SNPs (#sig. SNPs), and genomic inflation factor (GIF) are shown for the study site and years. The results of the conventional IGE model (Eq. S4) are shown on the right, which include LL, IGE weight 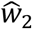, DGE-IGE covariance 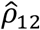, likelihood ratio *χ*^2^ and *p*-values over the model without *ρ*_12_. The site codes are shown as follows: Belgium, BEL; France, FRA; Italy, ITA; Poland, POL; Spain, ESP; and Switzerland, CHE. NA in the estimates means not available due to no phenotype records. The trait abbreviations are available in the caption of Figure 4. NA in the standard error (SE) means not applicable due to too small or large estimates.

Table S3. Variance component estimates and significant IGE SNPs in grape. The effective number of individuals with trait records (*N*), DGE variance 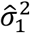, IGE variance 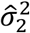, DGE-IGE covariance 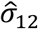, environmental variance 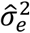, log-likelihood (LL), likelihood ratio and its p-values over a model without the covariance (*χ*^2^ and *p*), the number of significant IGE SNPs (#sig. SNPs), and genomic inflation factor (GIF) are shown for the three study years. The results of the conventional IGE model (Eq. S4) are shown on the right, which include LL, IGE weight 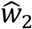, DGE-IGE covariance 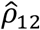, likelihood ratio *χ*^2^ and *p*-values over the model without *ρ*_12_. NA in the estimates means not available due to no phenotype records. The trait abbreviations are available in the caption of Figure 5. NA in the standard error (SE) means not applicable due to too small or large estimates. Bold rows highlight *p*<0.05.

## Supplementary Figures

**Figure S1.**
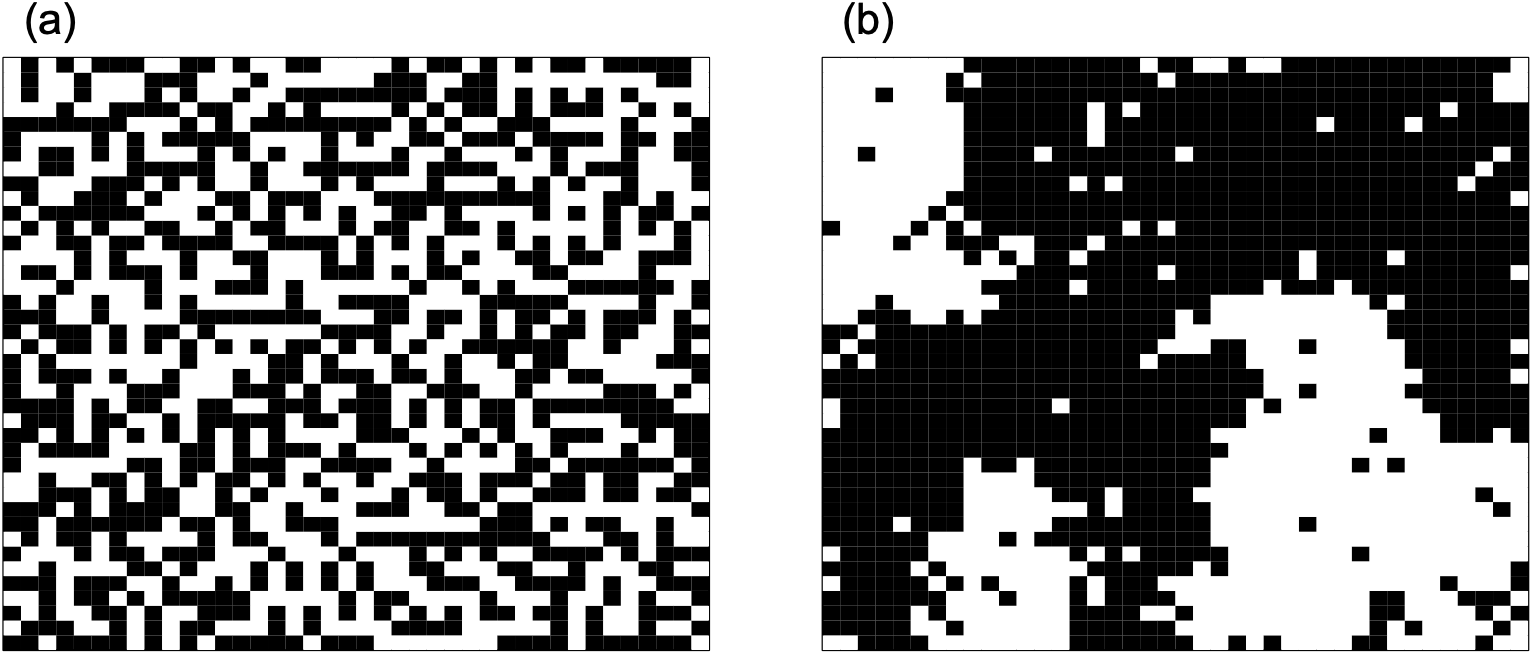
Ising model-based simulation of spatial patterns shaped by nearest neighbor interactions. The lattice space with 40 x 40 cells shows the outcome of 100-times update of the genotype value 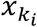 after a random configuration of two genotypes (black and white cells) following the fixed effects of Eq. S1.1, i.e., 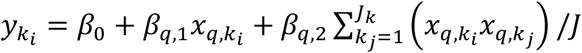. (a) The case of *β*_*q*,2_ = ™0.2, *β*_*q*,1_ = 0, *β*_0_ = 0. (b) The case of positive *β*_*q*,2_ = 0.2, *β*_*q*,1_ = 0, *β*_0_ = 0.

**Figure S2.**
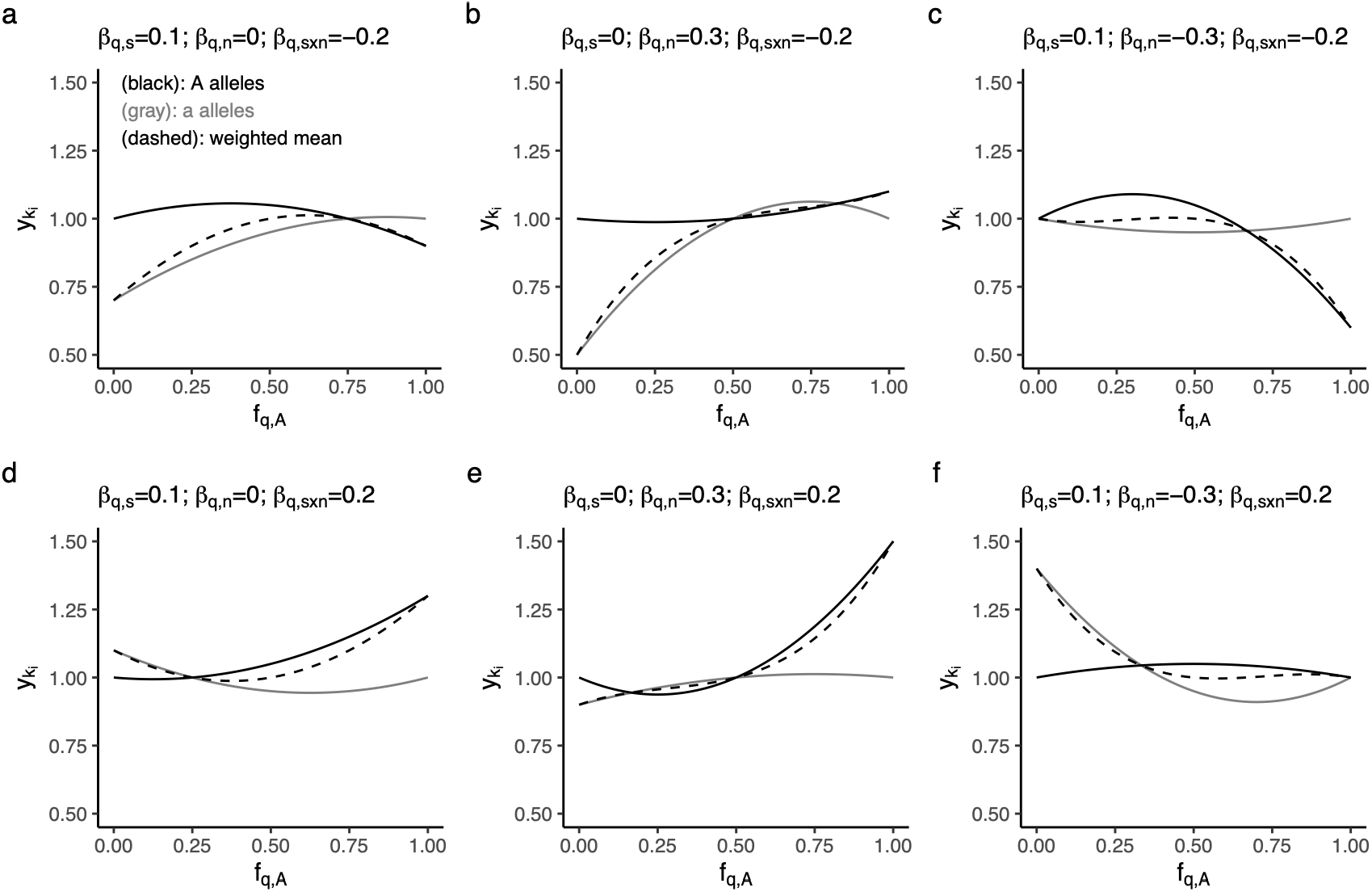
Numerical examples for the theoretical expectation between the phenotypic value 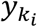 and the neighboring frequency of A alleles *f*_*q*,A_ at the *q*-th SNP locus within a local space *k*. Genotypes are encoded as {AA,Aa,aa} = {+1,0,-1} following Eq. S3.1. Black and grey lines indicate the phenotypic values of A and a alleles, respectively. Dashed curves represent a group mean defined by the weighted mean of the phenotypic values between A and a alleles. Panels (a-d) exemplify the respective cases of positive and/or negative *β*_*q*,n_(= *β*_*q*,12_) and *β*_*q*,s×n_(= *β*_*q*,2_) with zero or non-zero *β*_*q*,s_(= *β*_*q*,1_) and *β*_0_ = 1.0.

**Figure S3.**
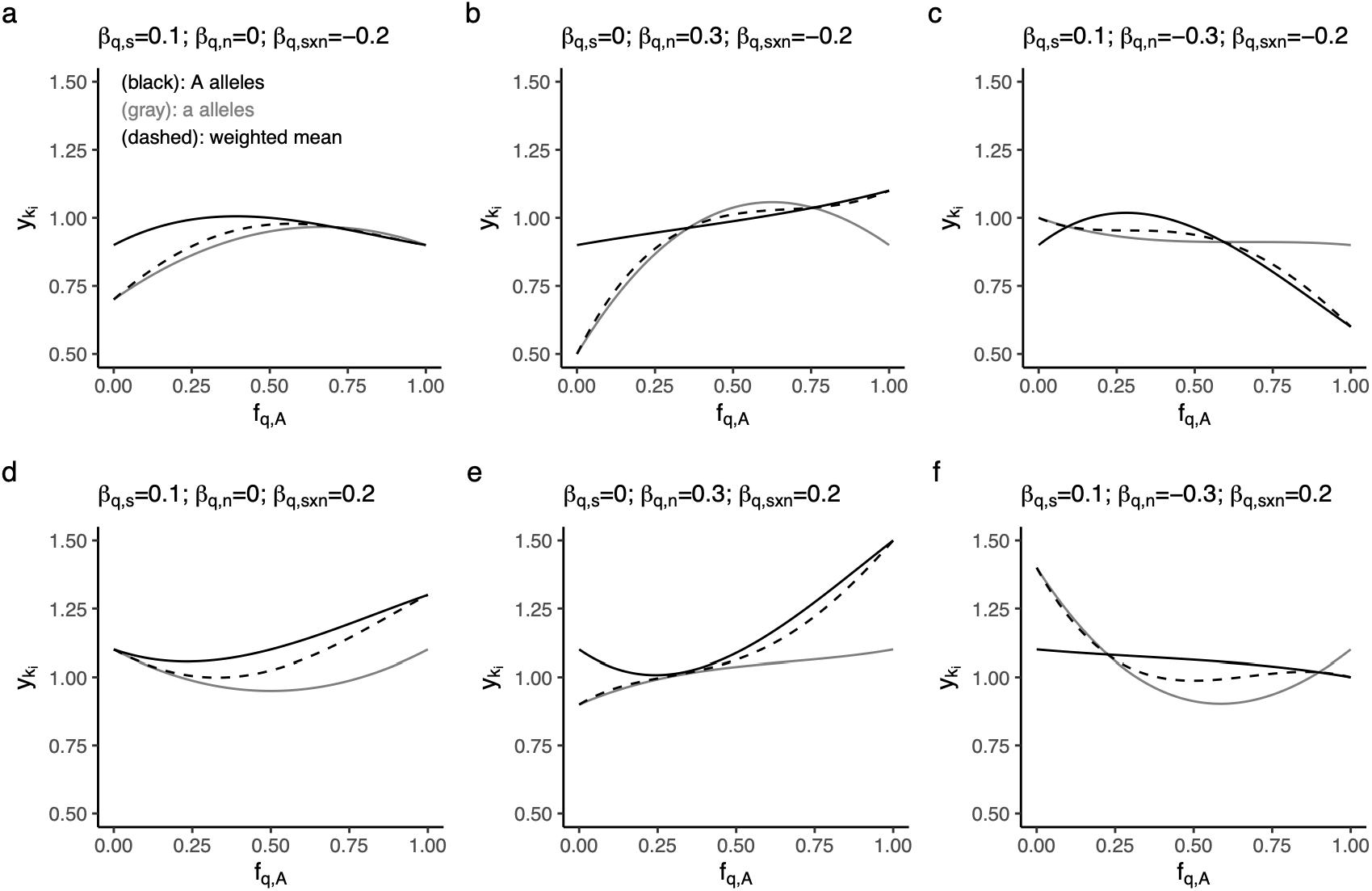
Numerical examples for the theoretical expectation between the phenotypic value 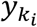 and the neighboring frequency of A alleles *f*_*q*,A_ at *q*-th SNP locus within a local space *k*. Self-genotypes are encoded as {AA,Aa,aa} = {+1,0,-1} but their interactions consider the half-similarity as *x*_*q*,AA_*x*_*q*,Aa_ = 0.5 and *x*_*q*,aa_*x*_*q*,Aa_ = ™0.5 following Eq. S3.2. Black and grey lines indicate the phenotypic values of A and a alleles, respectively. Dashed curves represent a group mean defined by the weighted mean of the phenotypic values between A and a alleles. Panels (a-d) exemplify the respective cases of positive and/or negative *β*_*q*,n_R= *β*_*q*,12_S and *β*_*q*,s×n_R= *β*_*q*,2_S with zero or non-zero *β*_*q*,s_R= *β*_*q*,1_S and *β*_0_ = 1.0.

**Figure S4.**
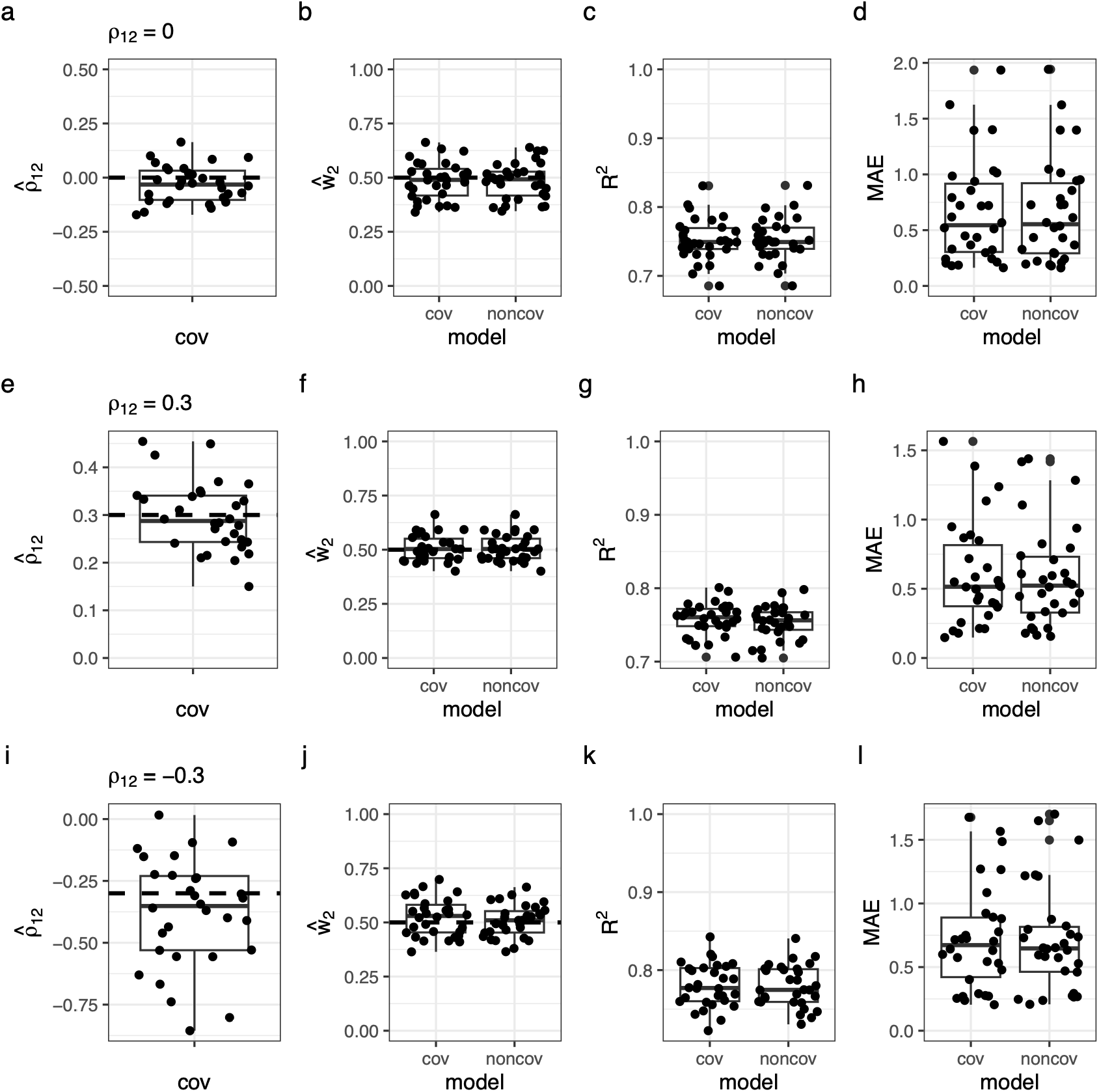
Variance components and genomic prediction of simulated data under the scenarios of moderate positive (*ρ* = 0.3: a-d), null (*ρ* = 0: e-h), or moderate negative (*ρ* = ™0.3: i-l) covariance between DGE and IGE. (a, e, and i) Estimated covariance 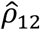 between polygenic DGE and IGE. Horizontal dashed lines indicate the true *ρ*_12_ value. (b, f, and j) Estimated weight of IGE 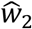 relative to that of DGE 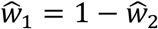. Models with or without the covariance 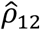 (cov or noncov) were fitted to simulated phenotypes. Horizontal dashed lines indicate the true *w*_2_ value. (c-d, g-h, and k-l) Genomic prediction based on the models with or without the covariance 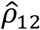 (cov or noncov). The model performance was evaluated using the coefficient of determination (*R*^2^: c, g, and k) and mean absolute error (MAE: d, h, and l) between the predicted phenotypic values **ŷ** and the observed trait values of the test dataset **y**. Box plots display the median and quartiles among 30 independent iterations, with whiskers extending to 1.5 × inter-quartiles.

**Figure S5.**
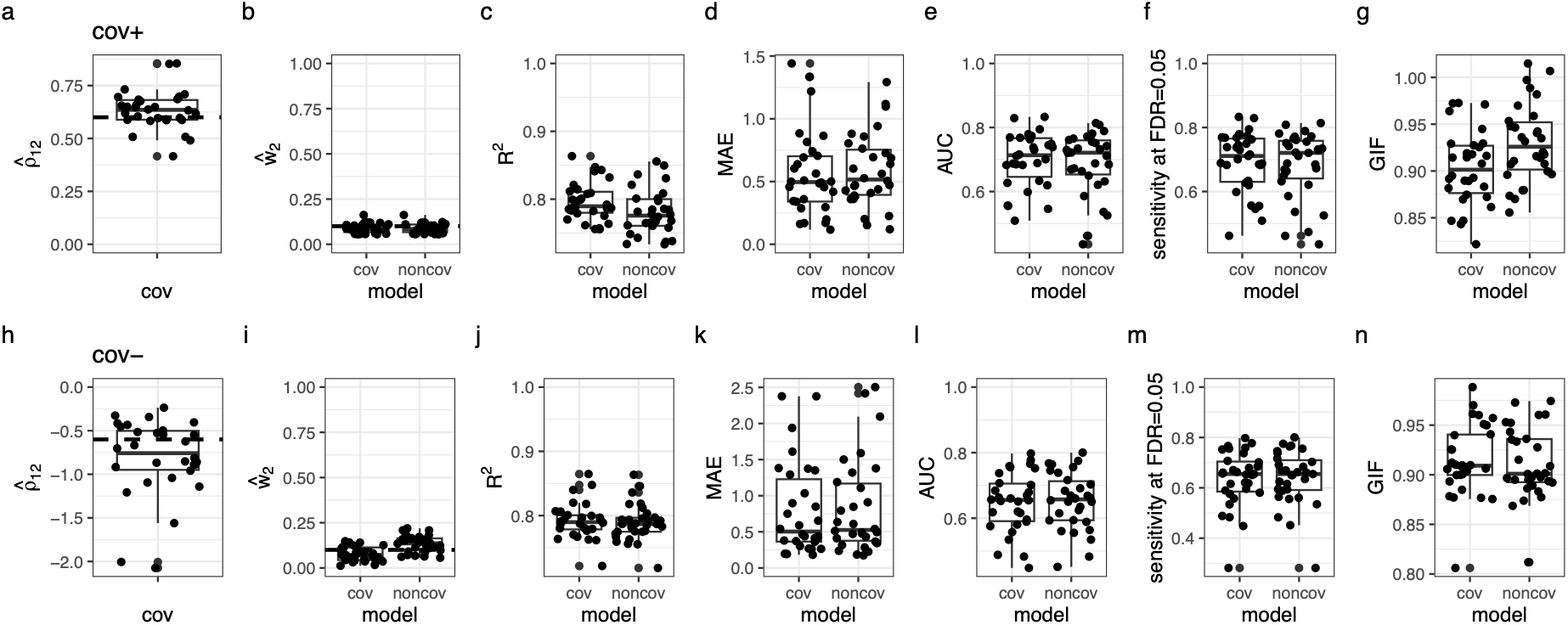
Benchmark simulations under the scenarios of positive or negative covariance with biased DGE-IGE weights as *w*_1_: *w*_2_ = 0.9: 0.1. The upper panels (a-g) show the results of positive covariance (*ρ* = 0.6) while the lower panels (h-n) show those of negative covariance (*ρ* = ™0.6). (a and h) Estimated covariance 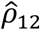 between polygenic DGE and IGE. Horizontal dashed lines indicate the true *ρ*_12_ value. (b and i) Estimated weight of IGE 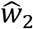 relative to that of DGE 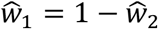. Models with or without the covariance 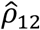 (cov or noncov) were fitted to simulated phenotypes. Horizontal dashed lines indicate the true *w*_2_ value. (c-d and j-k) Genomic prediction based on the models with or without the covariance 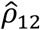 (cov or noncov). The model performance was evaluated using the coefficient of determination (*R*^2^: c and j) and mean absolute error (MAE: d and k) between the predicted phenotypic values **ŷ** and the observed phenotypic values of the test dataset **y**. (e-g and l-n) GWAS performance for the models with or without the covariance 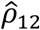 (cov or noncov) to detect causative IGE SNPs. The model performance was evaluated using the area under the ROC curve (AUC: e and l) and sensitivity at the stringent threshold at false discovery rate (FDR) = 0.05 (f and m). The genomic inflation factors (GIF: g and n) indicate the inflation (>1.0) or deflation (<1.0) of *p*-values compared to random expectation.

**Figure S6.**
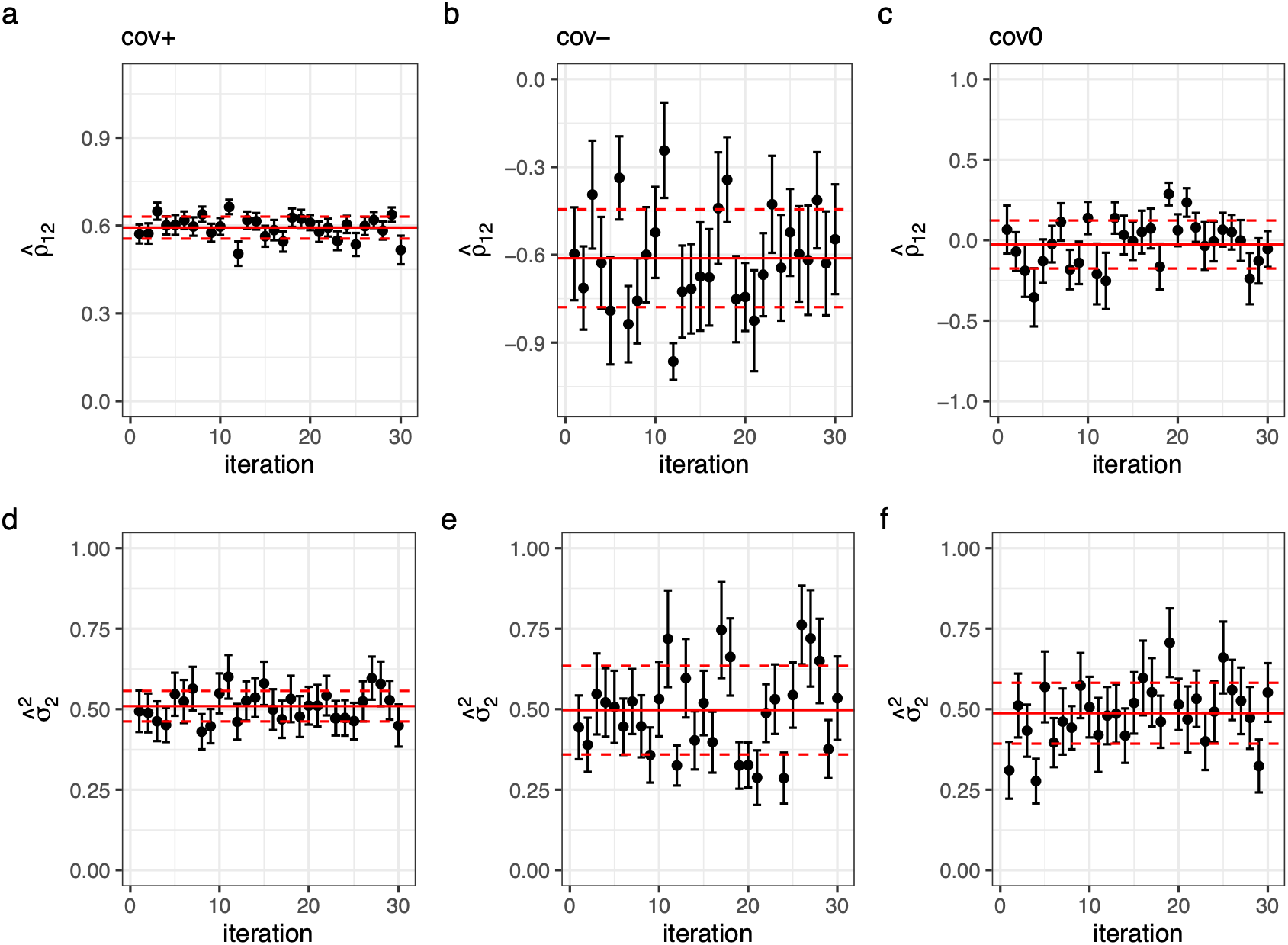
Standard errors (SEs) of the covariance parameter *ρ*_12_ and IGE variance 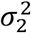 in the simulated data. Dot and error bars indicate the estimates ± SE. The horizontal red lines indicate the average (solid) ± standard deviation (dashed) of 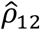 (a-c) or 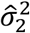 (e-f) over 30 iterations. The left (a and d), middle (b and e), and right (c-f) panels display the cases in which the true *ρ*_12_ >0, <0, and =0, respectively.

**Figure S7.**
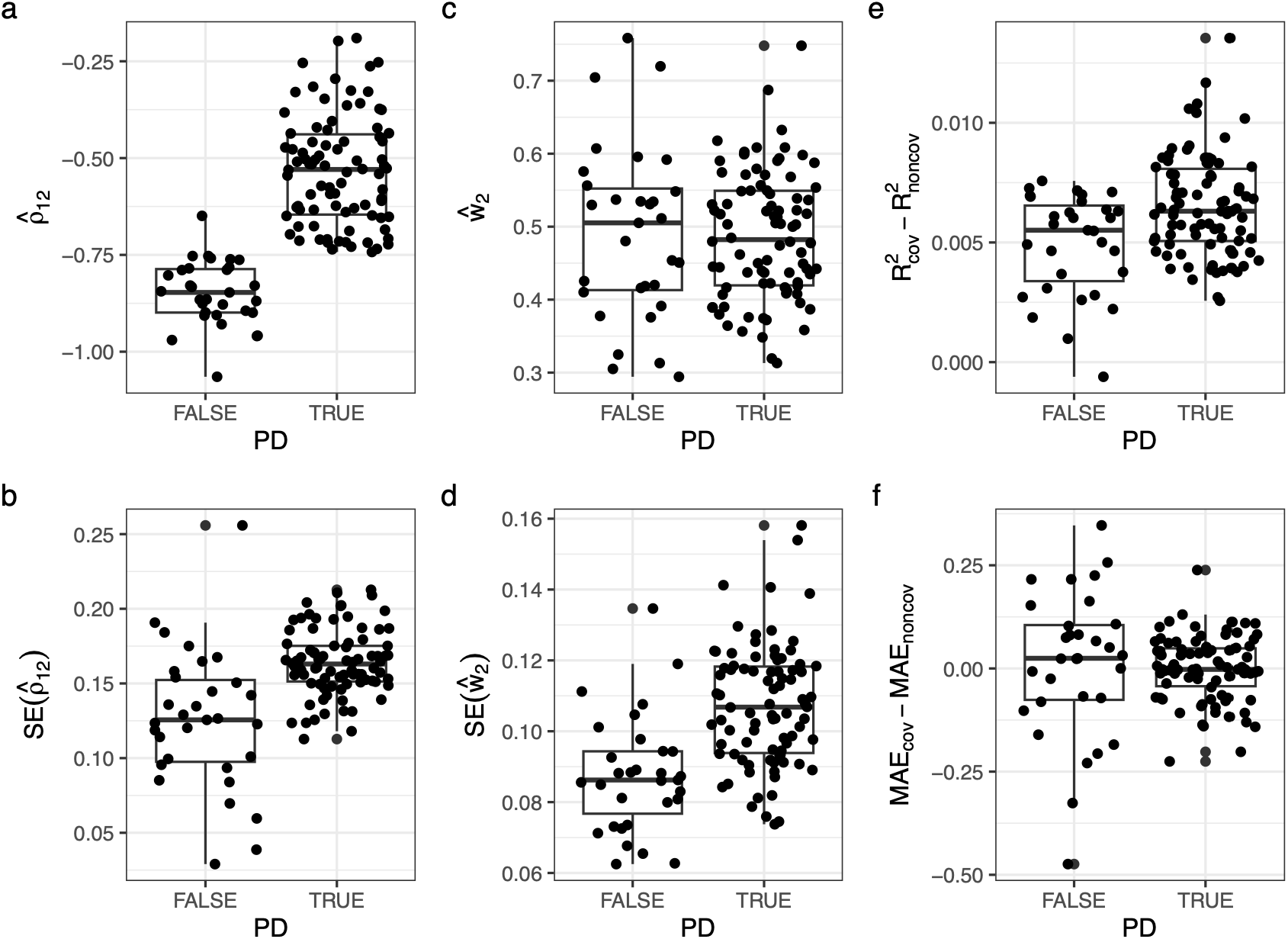
Influence of the nearest positive-definite approximation on parameter estimation and BLUP predictability. The x-axis displays the positive-definiteness (PD) of the weighted kernel **K**^′^ before approximation for the y-axes as follows. (a and b) Estimated covariance parameter 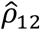 (a) and its standard error (b). (c and d) Estimated IGE weight 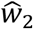 (c) and its standard error (d). (e and f) Difference of the coefficient of determination (*R*^2^) and mean absolute error (MAE) between the true and predicted BLUPs. The y-axes show *R*^2^ or MAE of the model with *ρ*_12_ ™ the model without *ρ*_12_, where zero indicates no difference between the models with or without *ρ*_12_. Simulations were performed 120 times with the true parameters of *ρ*_12_ = ™0.6, *w*_1_ = *w*_2_ = 0.5, and *V*_u_ = 0.6. Three iterations were discarded due to their convergence error.

**Figure S8.**
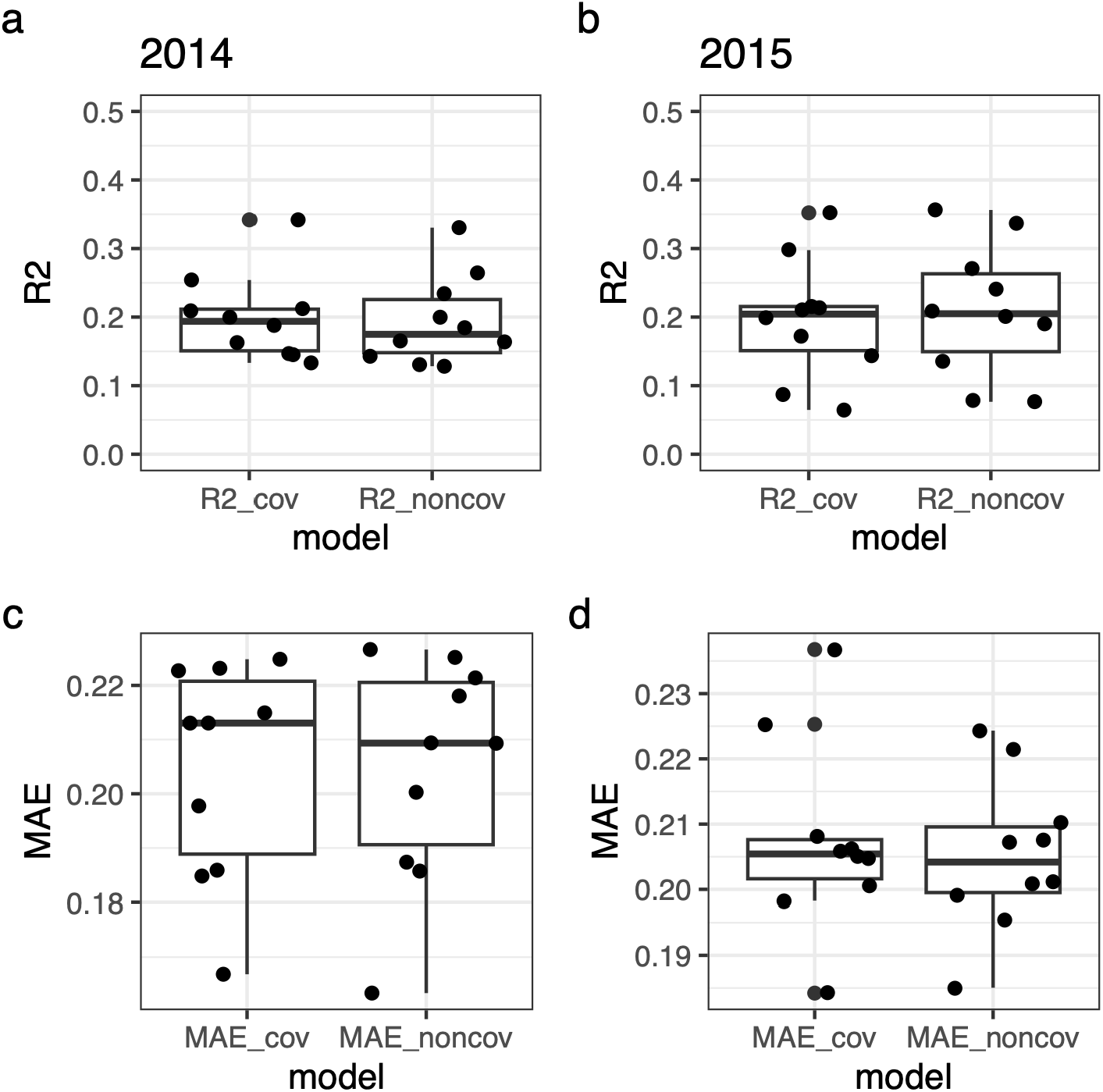
Genomic prediction of the basal area increment from 2014 to 2015 (BAI2014_2015) in aspen. The coefficient of determination (*R*^2^: a and b) and mean absolute error (MAE: c and d) of predicted phenotypic values **ŷ** among 10-fold cross-validations are shown for the data collected in 2014 (left) and 2015 (right).

**Figure S9.**
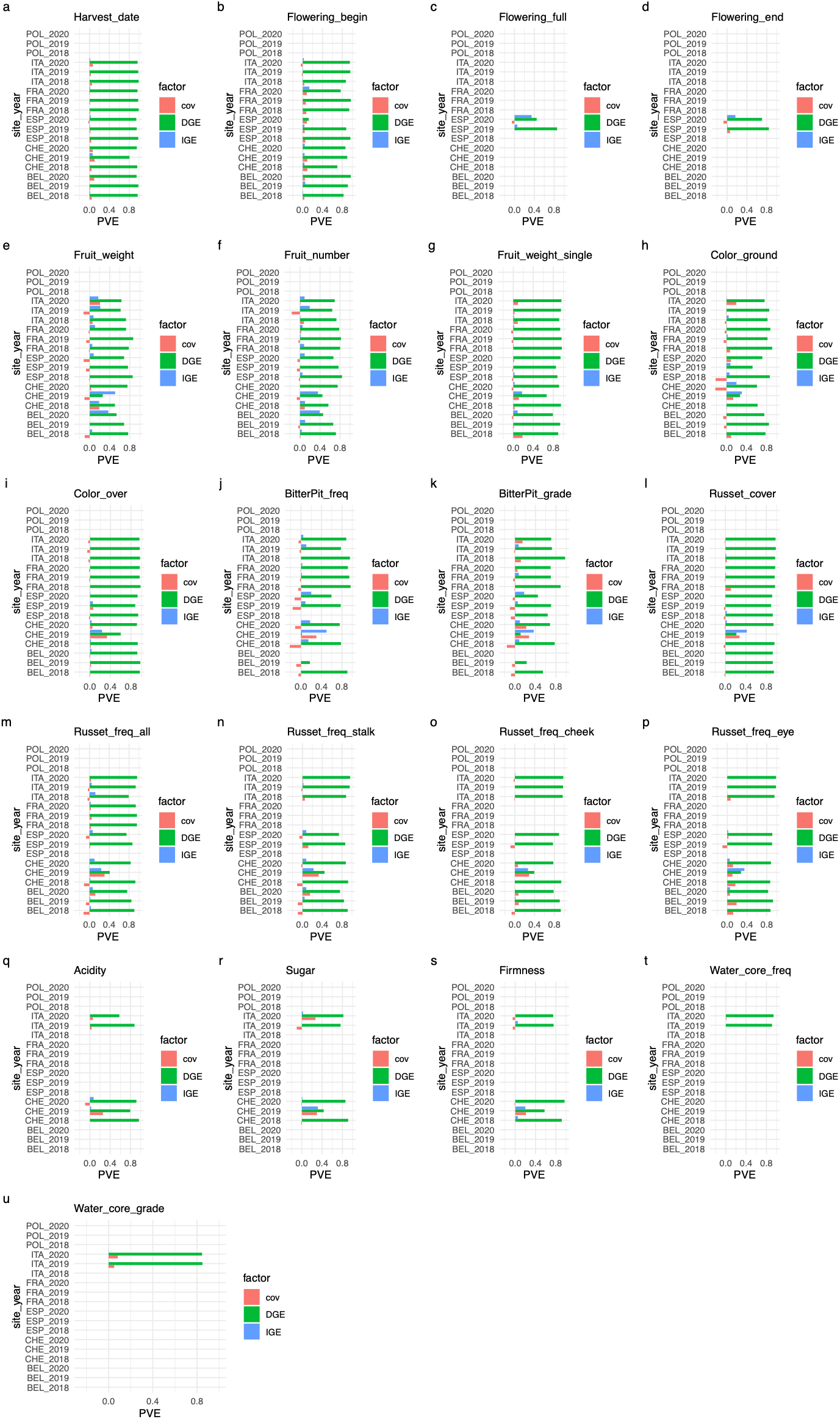
Proportion of phenotypic variation explained (PVE) by indirect genetic effects (IGEs), direct genetic effects (DGEs), and their covariance (cov) in 20 apple traits other than the flowering intensity, trunk diameter, and trunk increment given in the main text. Blue (upper), green (middle), and red (lower) bars indicate PVE by IGEs, DGEs, and their covariance for each trait, respectively. Blank bars indicate the absence of data.

**Figure S10.**
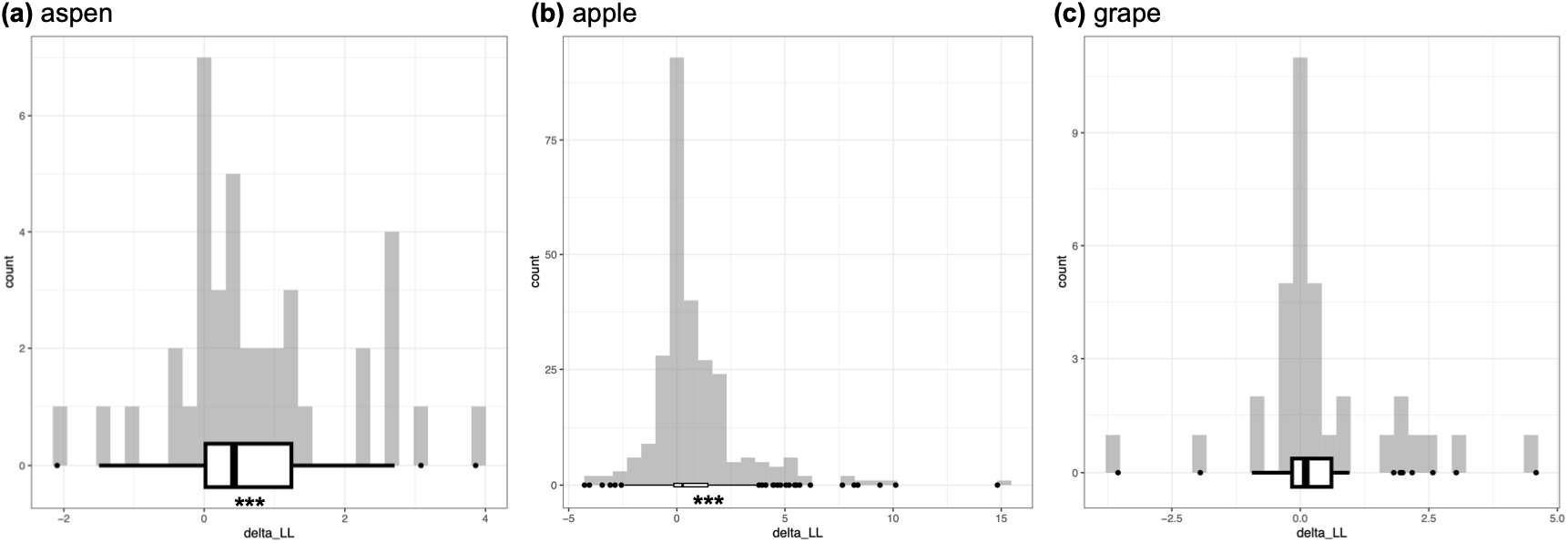
Difference of log-likelihoods between the proposed and conventional IGE model for the (a) aspen, (b) apple, and (c) grape data. The x-axes present the log-likelihood (LL) of our model (Eq. 1) - conventional IGE model (Eq. S4). Asterisks (^***^) indicate the significant deviation from zero with Wilcoxon signed rank tests at *p*<0.001. A single count corresponds to a trait per year/site (see Table S1, S2, and S3 for aspen, apple, and grape, respectively).

**Figure S11.**
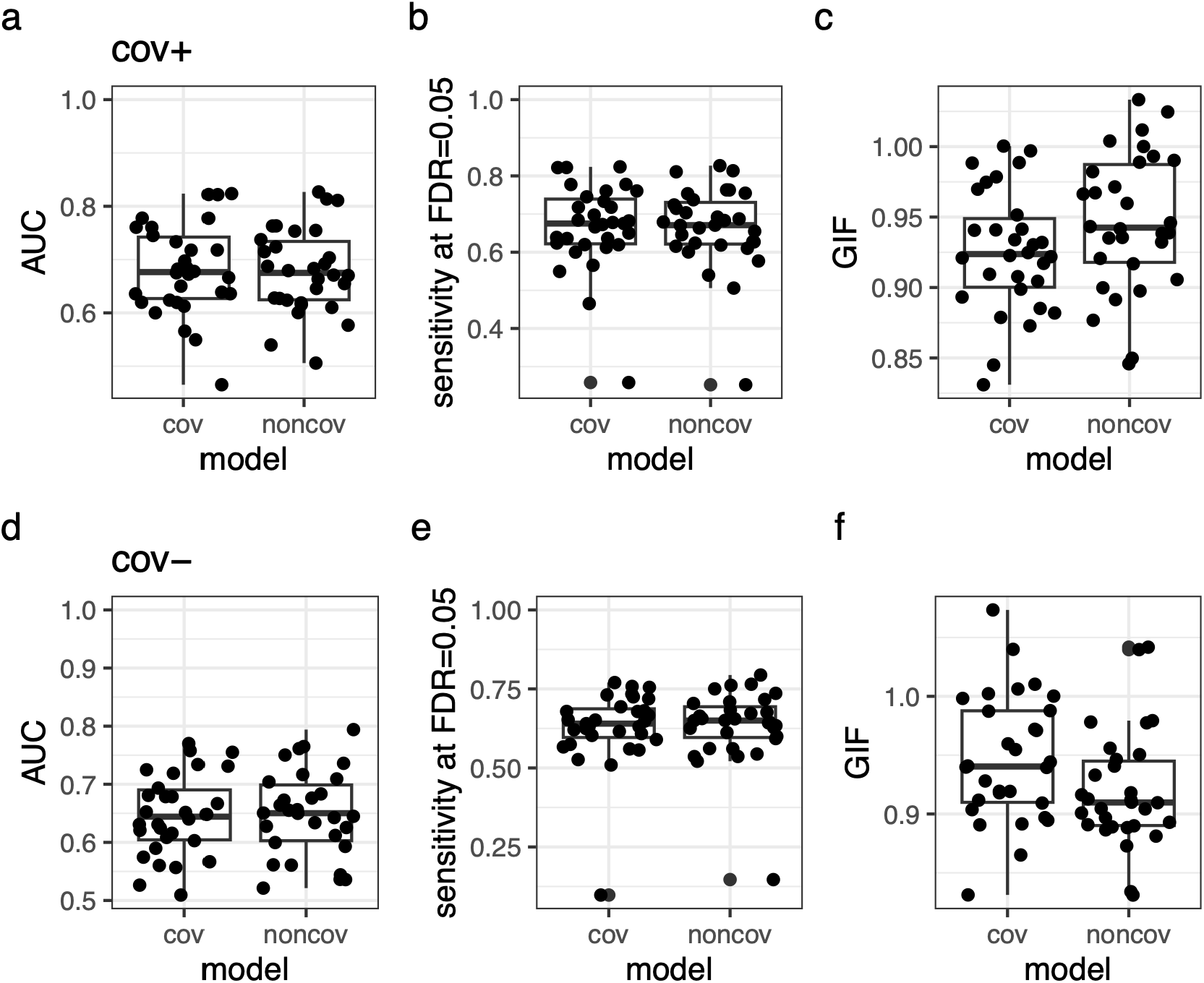
GWAS performance for the models with or without the covariance 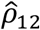 (cov or noncov) to detect causative IGE SNPs when IGE SNPs are uncorrelated with DGE SNPs. The other parameter settings were the same as those in Figure 2. The model performance was evaluated using the area under the ROC curve (AUC: a and d) and sensitivity at the stringent threshold at false discovery rate (FDR) = 0.05 (b and e). The genomic inflation factors (GIF: c and f) indicate the inflation (>1.0) or deflation (<1.0) of *p*-values compared to random expectation.

## Data accessibility

All source codes and input data are available in the GitHub (https://github.com/yassato/ngwas2) and Zenodo (https://doi.org/10.5281/zenodo.19362850) repositories. These repositories include a vignette that demonstrates how to implement relevant models using RAINBOWR, which can be viewed at https://yassato.github.io/ngwas2/. To make the present modeling easy, we provide an R package named ‘rNeighborLMM’ at GitHub (https://github.com/yassato/rNeighborLMM) and Zenodo (https://doi.org/10.5281/zenodo.19362843).

## Author contributions

**YS**: Conceptualization, Methodology, Software, Data curation, Investigation, Formal analysis, Validation, Visualization, Funding acquisition, Writing - Original Draft; **KH**: Conceptualization, Methodology, Software, Funding acquisition, Writing - Review & Editing.

## Acknowledgements

We thank M. Jung and H. Muranty for providing complete genotype data of the apple REFPOP project prior to publication; H. Barker and J. Riehl for addressing our questions about the aspen field arrangement and data annotation; and H. Iwata, S.E. Wuest, and M. Roth for discussions during the study. We acknowledge that the computing resource was provided by Human Genome Center at the University of Tokyo, Japan (http://sc.hgc.jp/shirokane.html). We used Paperpal by Editage (https://paperpal.com/) for language corrections but did not use any generative AI to draft the manuscript.

